# Rapid, accurate long- and short-read mapping to large pangenome graphs with vg Giraffe

**DOI:** 10.1101/2025.09.29.678807

**Authors:** Xian Chang, Adam M. Novak, Jordan M. Eizenga, Jouni Sirén, Jean Monlong, Shloka Negi, Francesco Andreace, Sagorika Nag, Konstantinos Kyriakidis, Glenn Hickey, Stephen Hwang, Emmanuèle C. Délot, Andrew Carroll, Kishwar Shafin, Pi-Chuan Chang, Faith Okamoto, Benedict Paten, the Human Pangenome Reference Consortium

**Author notes:** these authors contributed equally to this work (Xian Chang, Adam M. Novak).

## Abstract

We previously introduced Giraffe, a short-read-to-pangenome graph mapper available in the vg pangenomics toolkit. Giraffe was fast and accurate for mapping short reads to human-scale pangenomes, but struggled with long reads. Long reads present a unique challenge to pangenome mapping algorithms due to their length and error profile, which allow them to take more topologically complex paths through the pangenome graph and increase the possible search space for the algorithm. We present updates to Giraffe that allow it to quickly and accurately map long reads to pangenome graphs. For both short and long reads, Giraffe mapping to a pangenome containing data from more than 450 human haplotypes, generated by the Human Pangenome Reference Consortium, is comparable in speed to linear mappers to human reference genomes; Giraffe is also over an order of magnitude faster than GraphAligner, the current state-of-the-art long-read-to-pangenome mapper. Its alignments produce similar or improved small and structural variant calling results, compared to those from commonly used graph-based and linear mappers. We additionally demonstrate using Giraffe’s long read alignments in a pangenome-guided assembly workflow, which is capable of producing more contiguous local assemblies than Hifiasm in our test regions.

## 2 Introduction

Pangenomes are emerging as a promising alternative to the standard single linear reference genome, because they are capable of representing the genetic diversity present in a population [1, 2]. There have been several recent efforts to establish human reference pangenomes, either targeted (at particular countries [3], ancestries [4], or sets of ethnicities [5]), or global across the human population [6]. Pangenomes are commonly represented as variation graphs, where nodes represent nucleotide sequences, edges represent adjacencies between sequences, and paths through the graph represent the original haplotype sequences of the pangenome [7]. These graph structures are an efficient representation of pangenomes, but the complex topology of graphs makes them more difficult to work with than linear reference genomes, especially as the graphs grow in sequence content and genetic diversity. In particular, mapping sequencing reads, a critical first step for many common bioinformatics pipelines, requires specialized data structures and algorithms to deal efficiently with a graph reference.

Read mappers that align to a standard linear reference [8, 9, 10] commonly use a **seed-and-extend** approach [11]. Short **seed** alignments are found using an index of the reference genome. Many algorithms then partition seeds into groups that are likely to have come from the same alignment to the reference. Seeds are then **extended** into the full base-level alignment using dynamic programming.

Existing graph mappers use versions of these basic components that have been adapted to a graph context. Early graph alignment algorithms were designed to align reads to partial order alignment (POA) graphs that themselves represent multiple sequence alignments [12]. These POA aligners generalized standard dynamic programming algorithms to align to acyclic graphs [12, 13]. The original `vg map` algorithm further extended the POA algorithm to cyclic graphs by “unrolling” the graph to remove cycles, then applying POA to implement its extension step [7]. GraphAligner, currently the only practical tool for mapping long reads to arbitrary graphs, generalizes the linear *Shift-And* exact string matching algorithm [14] for its seeding step, and Myers’ bitvector alignment algorithm [15] for its extension step, to align to cyclic graphs [16].

The original short read Giraffe relied on embedded haplotype sequences to map to the graph [17]. This set of embedded haplotypes could first be **personalized** using **haplotype sampling**, to remove local haplotypes not relevant to a particular sample [18]. Next, seeds were found by using a minimizer index over the haplotypes, and clustered by using a distance index to find minimum graph distances. Seeds within promising clusters were gaplessly extended along the haplotypes, and finally the extension was completed using dynamic programming against the graph. Due to the fast gapless extension step, in which the entire alignment was usually found, Giraffe was very fast for short reads [17]. However, being designed for short reads, where gaps are rare and a read will usually follow a single known haplotype for its entire length, Giraffe quickly overwhelmed our computational resources when applied to long reads.

Long reads are another rapidly growing technology with great potential to improve genomic analyses. Currently, High Fidelity (HiFi) reads from Pacific Biosciences have lengths between 10 and 25 kbp and accuracy of up to 99.95% [19, 20]. The most recent R10 chemistry from Oxford Nanopore Technologies achieves read lengths on the order of 10-100 kbp, with ultra-long reads achieving lengths up to 882 kbp [21], and accuracy of over 99% [22, 23]. Because of their increased length, these technologies are useful for resolving complex and large structural variants, and for identifying long-range linkages between alleles. Both HiFi and R10 sequencing technologies have gained widespread use in large-scale genomics projects [24, 25] due to this improved resolution ability, as well as their lowering cost and high throughput.

Despite the growing popularity of both pangenomes and long reads, there are currently few tools for mapping long reads to pangenomes. The length and error profile of the reads makes every stage of a mapping algorithm more difficult. Gapped alignment is a particularly challenging problem with long reads: even in a linear context, long reads can produce dynamic programming matrices that are unmanageably large. The problem becomes even harder in a graph context, as aligners must also navigate potentially large and complex graph topologies. To deal with the increased difficulty of mapping long reads, many tools [10, 26, 27, 28] build on the seed-and-extend approach by adopting a **seed-and-chain** or **seed-chain-extend** strategy [11]. After seeds are found, and before they are extended into alignments, **chains** of co-linear seeds are identified. Then, extension consists of smaller alignment problems between consecutive seeds in a chain.

Chaining requires an ordering and a distance metric between seeds, both of which are ambiguous and more expensive to compute in a graph. Chaining is particularly difficult in graphs that contain cycles, as a node may occur before or after itself or any other node in the cycle [29, 28, 30].

Existing graph chaining algorithms use distance calculations based on linear approximations of the graph. Some tools [29, 28, 31] are based on an algorithm from Mäkinen et al. [30] that uses a minimum path cover of the graph to determine reachability and distance. Other tools might enumerate shortest-length walks between graph positions (Minigraph [26]) or estimate linear distances based on chains in the snarl decomposition (GraphAligner [32]).

As a result of these different heuristics for overcoming the challenges of long read-to-pangenome mapping, different mappers have different advantages and limitations.

GraphAligner, for example, can accurately map reads to arbitrary graphs, but it is slow, particularly as the complexity of the graph increases [32]. Minigraph [26], which was originally designed to map assemblies to graphs containing only structural variants (SVs), struggles to map to graphs containing small insertions or deletions (indels) or single nucleotide variants (SNVs). PanAligner [27] uses its own graph chaining algorithm with components from Minigraph, and GraphChainer [28] uses its own graph chaining algorithm with components from GraphAligner. Minichain uses a haplotype-aware chaining algorithm that prioritizes alignments that remain on the same haplotype [31]. PanAligner has the same limitations as Minigraph and was an order of magnitude slower than GraphAligner in our initial tests. GraphChainer and Minichain can only map to graphs without cycles.

We present an updated version of Giraffe capable of mapping both long and short reads. The long read Giraffe algorithm uses a seed-chain-extend framework, using a novel data structure for computing distances for chaining. We show that Giraffe is among the fastest graph-based or linear read mappers, while still being competitive in mapping accuracy and downstream variant calling results. Finally, we use Giraffe’s long read mappings in a **haplotype-resolved pangenome-guided assembly** workflow for assembling a specific region of interest, solving a relaxed version of the **haplotype-resolved *de novo* assembly** problem addressed by tools like Hifiasm [33].

## 3 Results

### 3.1 Long read Giraffe algorithm overview

The long read Giraffe algorithm is adapted from the original short read Giraffe algorithm [17] to efficiently deal with the longer sequence and higher error rate of long reads (Figure 1). The seed-and-extend short read algorithm has been expanded into a seed-chain-extend algorithm of the kind typical of long read mappers [11], with additional improvements that are useful in a long read context. For seeding, the long read algorithm uses weighted minimizer seeds, which prioritizes seeds with fewer hits in the graph (Figure 1 C). For chaining, we address the challenges of complex graph topology using a novel data structure for finding graph-distances between seeds that are supported by the corresponding read-distances (Figure 1 D). Using this data structure, the algorithm computes co-linear chains of seeds that occur in the same order in the read and graph, which form the backbone of the final alignment (Figure 1 E). Finally, for extension, base-level alignment is done in between consecutive seeds in the chain using alignment algorithms that limit the problem to alignment either against linear haplotypes or within smaller bands of the alignment matrix (Figure 1 F).

**Figure 1:**
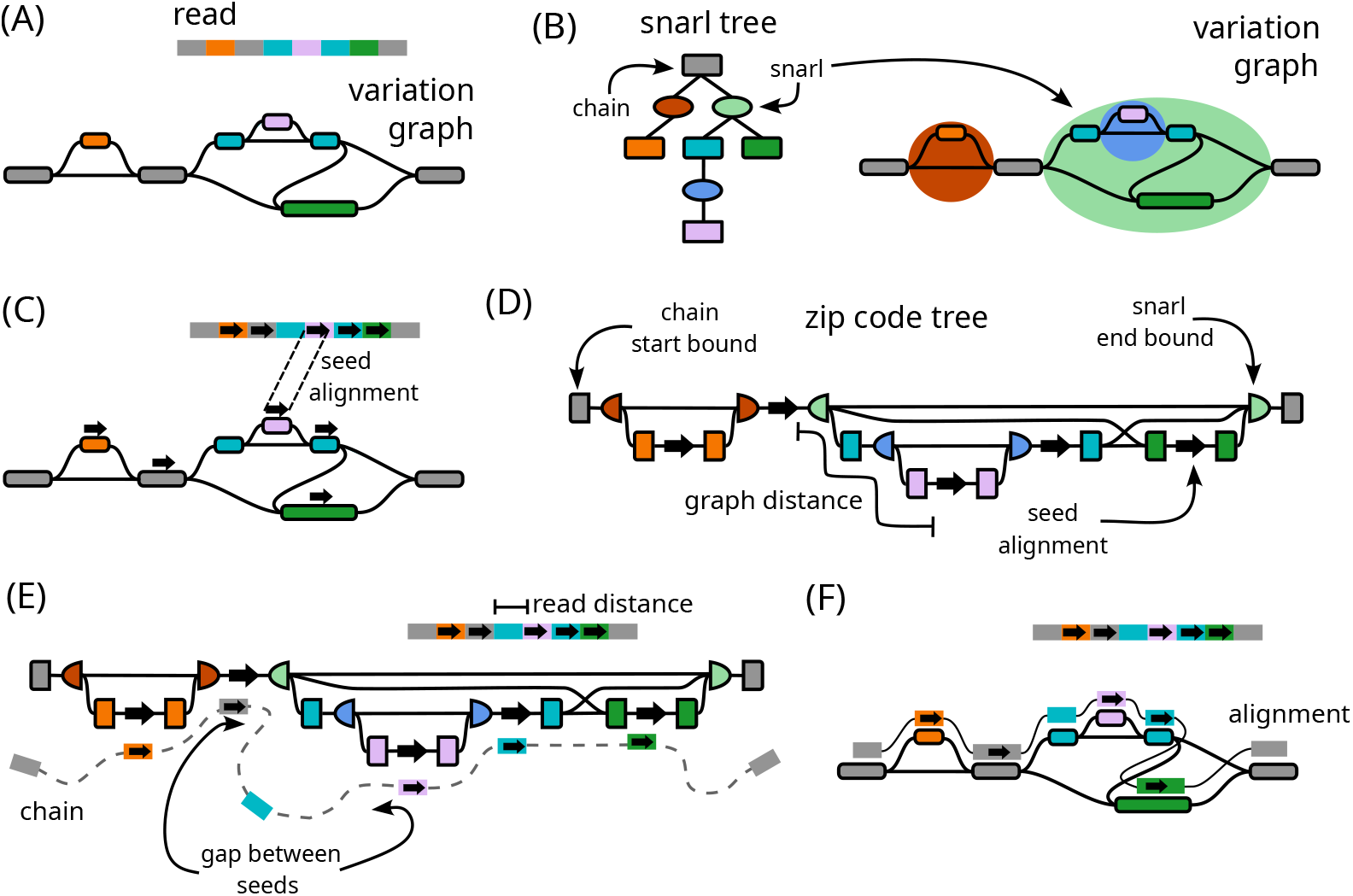
Overview of the long read Giraffe algorithm. (A) The inputs to the algorithm: the query read and the reference variation graph. (B) The variation graph with its corresponding snarl tree. Rectangles in the snarl tree represent chains and ovals represent snarls. Chains on the snarl tree are colored according to their node children. Nodes are not shown on the snarl tree. (C) Seeds are found using a minimizer index. Arrows on the read and reference graph represent seed alignments. (D) A zip code tree is constructed to represent the connectivity between seeds on the graph using snarl tree relationships. Nodes in the zip code tree represent either a seed on the graph, or the boundary of a snarl or chain containing a seed. The nodes are ordered according to a pre-and-post-order traversal of the snarl tree. Edges connect adjacent seeds and boundaries, and are labeled with minimum distances from the variation graph. Distances between seeds in the graph can be calculated in a traversal of the zip code tree. (E) Co-linear chaining is done by dynamic programming over the seeds to maximize the coverage over the read and minimize the gap cost of consecutive seeds. The cost of a gap between seeds is found using read distances from the read and graph distances from a traversal of the zip code tree. (F) Base-level alignment is done between adjacent seeds in a chain, and out from the first and last seeds to produce the full-length alignment.

### 3.2 Experiment setup: graphs, reads, and mappers

We compared Giraffe to a variety of linear mappers aligning to the CHM13 reference genome, and to graph mappers aligning to graphs constructed from the Human Pangenome Reference Consortium’s (HPRC) release 2 assemblies (Section S5.1). We used the HPRC graph constructed with just Minigraph (HPRC-Minigraph) as well as the graph constructed with the Minigraph-Cactus pipeline with allele frequency filtering (HPRC-d46) [6]. In a previous study [18], we found that short read Giraffe performed better when using personalized pangenome graphs (see 2). We therefore tested Giraffe on graphs that were haplotype-sampled to contain only 16 haplotypes and whichever of the two linear references was appropriate (HPRC-Sampled16). The haplotype sampled graph used a base graph constructed with the Minigraph-Cactus pipeline with selective clipping (HPRC-clipped). Our haplotype-sampling pipeline requires k-mers from a whole-genome read set, so we generated a different sampled graph for each sample and sequencing technology, by using an appropriate set of real reads (Section S5.1). Finally, we also used Giraffe to map to a negative control variation graph containing only CHM13 (also referred to as CHM13).

We mapped real datasets of long read technologies (HiFi and R10) and short read technologies (Illumina and Element) reads (Section S5.2).

We compared Giraffe to Minimap2 [10], Winnowmap [34], GraphAligner [16, 32], and Minigraph [26] for mapping HiFi and R10 reads; PBMM2 [10, 35] for mapping HiFi reads; and to Minimap2 and BWA-MEM [9] for mapping Illumina and Element reads (Section S5.3). Giraffe was run using the long read algorithm on HiFi and R10 reads, and using the paired-end short read algorithm for Illumina and Element reads. GraphAligner was run using the default parameters on simulated reads and when aligning to the HPRC-Minigraph graph with R10 reads, but we were unable to finish mapping real HiFi or R10 reads to the HPRC-d46 graph, or real HiFi reads to the HPRC-Minigraph graph, within the seven-day time limit of our servers, so this was done using a parameter set recommended by the developer to optimize for speed, which we call the “fast” parameters (Section S5.3).

### 3.3 Mapping evaluation

We used simulated read datasets to evaluate mapper performance on HiFi, R10, Illumina, and Element reads. Each dataset contained 1 million reads simulated to match the error profiles of the real datasets (Section S5.2).

Giraffe’s accuracy was similar to that of other mappers, and it had the highest true positive rate for HiFi reads when mapping to the allele frequency-filtered HPRC-d46 graph (Figure 2, Tables S1-S4). Giraffe generally had better mapping quality calibration than other mappers (Figure S4) and the fewest incorrectly mapped reads with a mapping quality (MAPQ) of 60, meaning that it was less likely to overestimate its accuracy. (Tables S1-S4).

**Figure 2:**
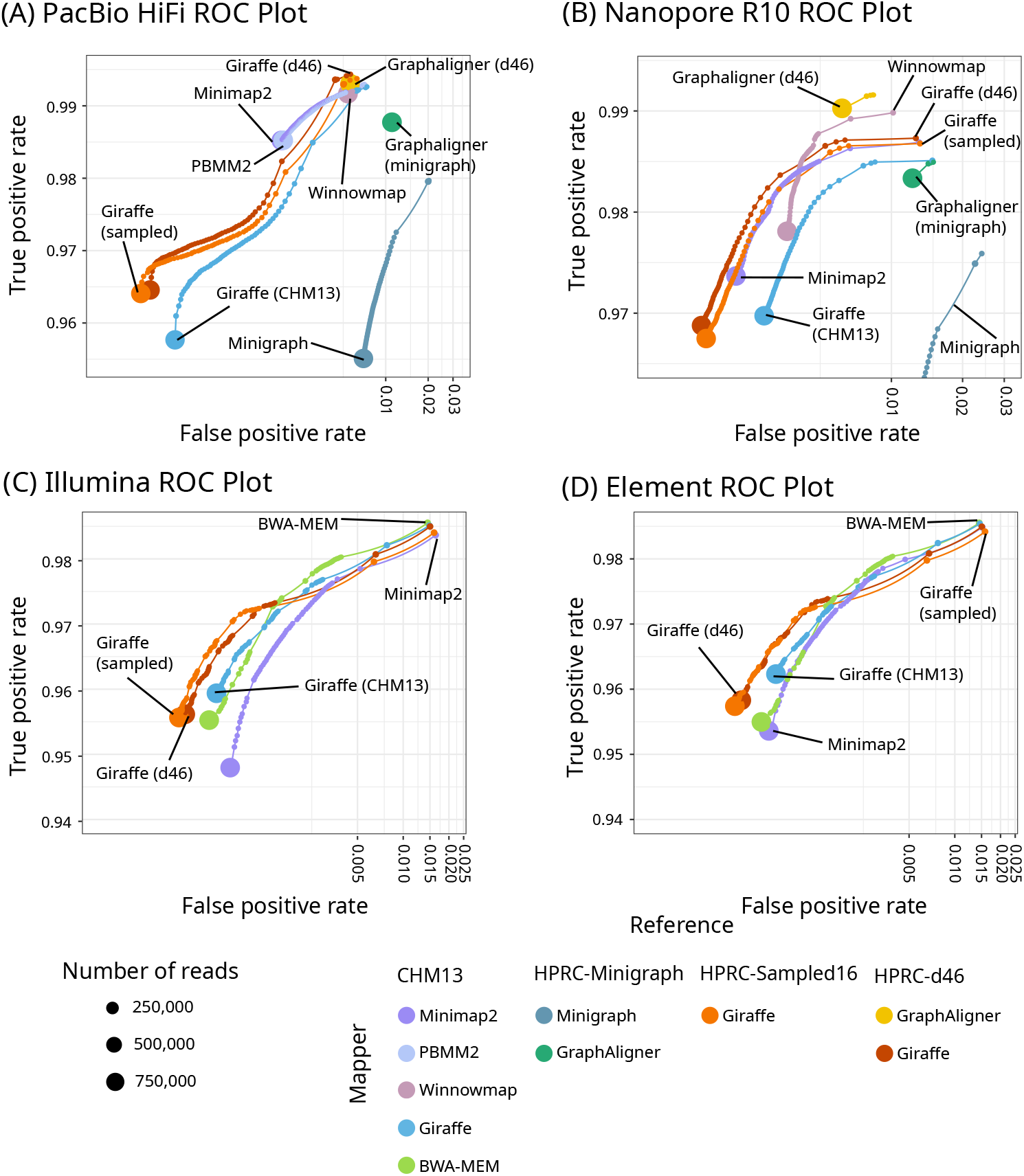
Mapping accuracy on simulated reads. ROC plots for one million reads simulated to resemble HiFi (A), R10 (B), Illumina (C), and Element (D) reads, zoomed in to show the separation of the curves. The results are stratified by mapping quality; the size of each point represents the log-scaled number of reads with the given mapping quality. X-axes are log-scaled. Linear mappers (Minimap2, PBMM2, Winnowmap, BWA-MEM) were used to align to CHM13, Giraffe to the linear CHM13 graph, and graph mappers (Giraffe, GraphAligner, Minigraph) to the HPRC-d46 graph (Giraffe and GraphAligner), HPRC-Minigraph graph (Minigraph), and the HPRC-Sampled16 graph (Giraffe).

We also mapped real read sets for each sequencing technology (Section S5.2), to evaluate unaligned bases, and to assess speed and memory use. Because GraphAligner exceeded the seven-day time limit of our servers in some cases, we re-ran it using a parameter set recommended by the developer (Section S5.3). We evaluated the base-level alignments of real long reads by comparing the total number of different types of unaligned bases (soft-clipped, hard-clipped, and unmapped) that each mapper produced; for short reads, we looked at the fraction of unmapped bases (Figure 3). For long reads, again, results were similar between Giraffe and most other mappers. For short reads, we found that the existing short read Giraffe algorithm could leave noticeably more reads unmapped than other mappers. We noted the same pattern of unmapped reads when mapping simulated reads. Of the simulated reads that Giraffe left unmapped, all were incorrectly mapped by BWA-MEM and Minimap2.

**Figure 3:**
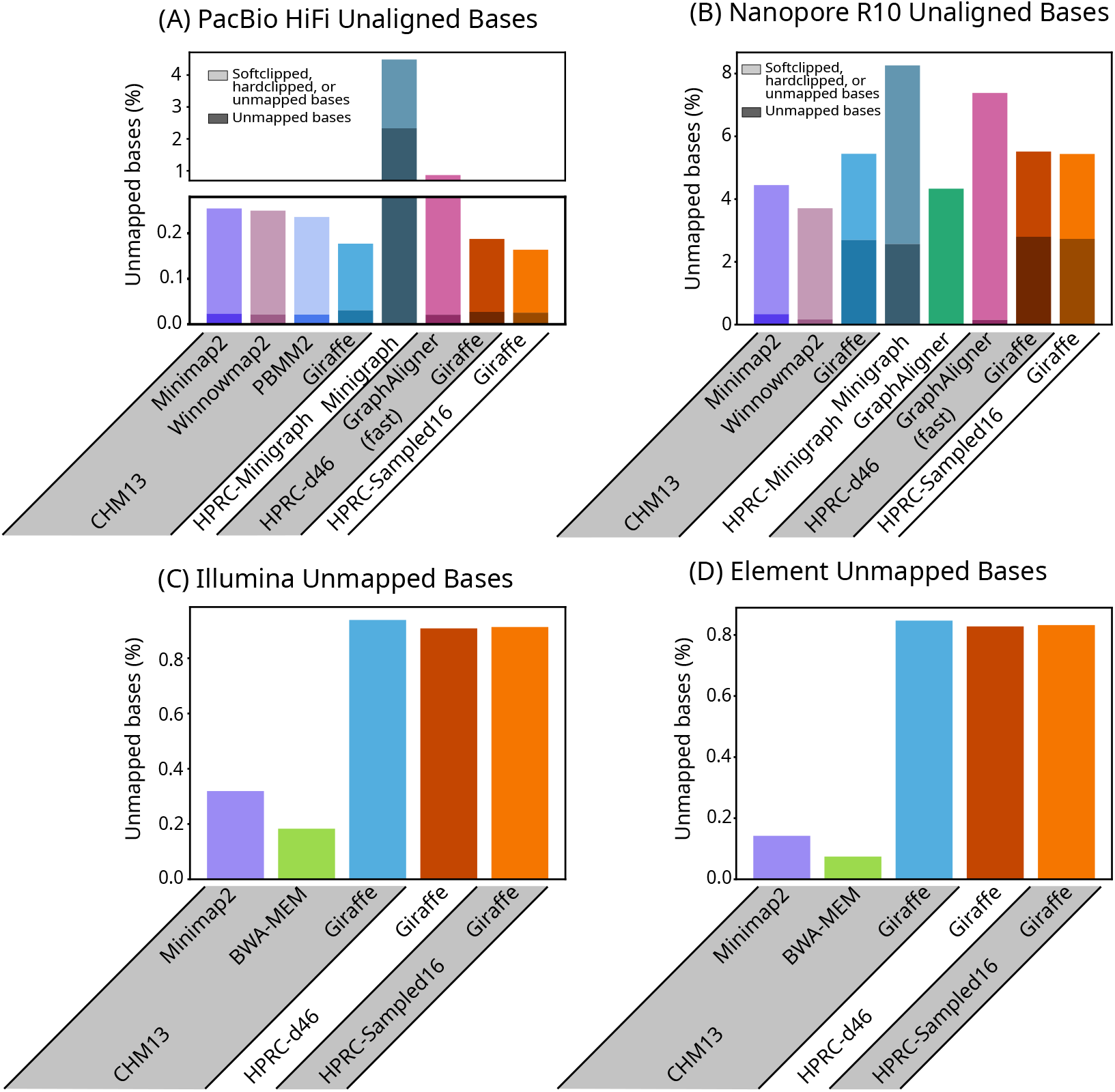
Unaligned bases. Bar plots of the percent of base pairs that were unaligned due to being softclipped, hardclipped, or in reads that were left unmapped, in real HiFi (A) and R10 (B) reads, and the percent of bases that were unmapped in Illumina (C) and Element (D) reads. Bases that were unaligned due to being in unmapped reads are represented by the lower, darker portion of each bar in (A) and (B), and are the entirety of each bar in (C) and (D).

Giraffe was about an order of magnitude faster than GraphAligner’s fast mode (Figure 4). Giraffe’s speed and memory use varied with the complexity of the reference graph it aligned to, with the simplest CHM13 graph being the fastest and lowest memory and the most complex HPRC graphs (HPRC-d46 and HPRC-Sampled16) being the slowest with the biggest memory footprint (Figure 4).

**Figure 4:**
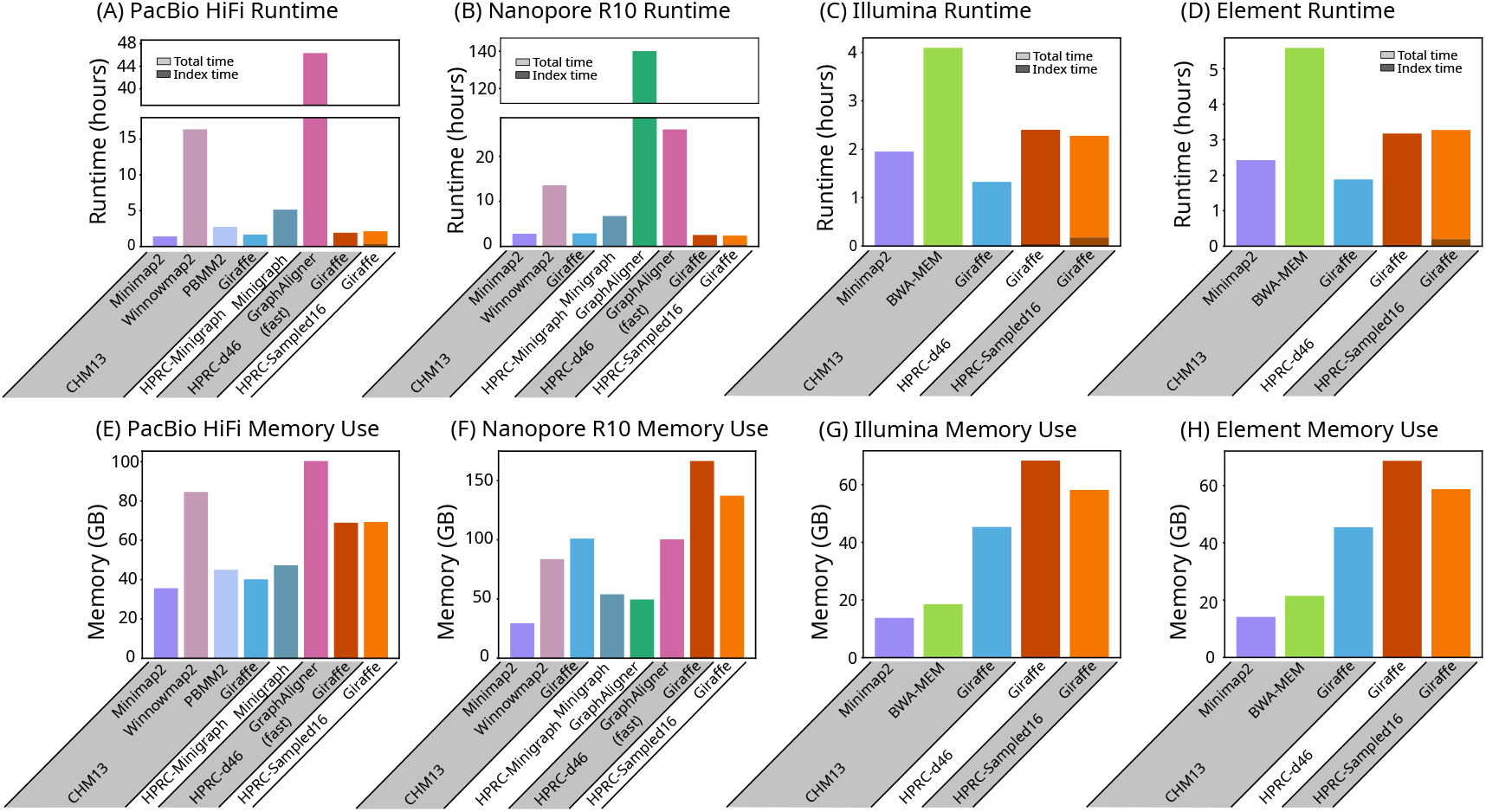
Runtime and memory use. Runtime (A, B, C, and D) and memory use (E, F, G, and H) of each mapper on real PacBio HiFi (A and E), Oxford Nanopore R10 (B and F), Illumina (C and G), and Element (D and H) reads. The lower, darker portion of each runtime bar shows the part of the runtime used for loading and building indexes, as reported by each mapper, including the time for haplotype sampling for Giraffe. * GraphAligner exceeded the 7-day time limit of our servers when run with the default settings on HiFi and R10 reads on the HPRC-d46 graph, and on HiFi reads on the HPRC-Minigraph graph. It was re-run using a parameter set (GraphAligner-fast) that was provided by the developer (Section S5.3).

### 3.2 Small variant calling with DeepVariant

To more robustly assess the quality of Giraffe’s long read alignments, we compared the performance of DeepVariant [36] using reads mapped with Giraffe and Minimap2. For Giraffe, PacBio and R10 reads from the HPRC (for HG002) were mapped to the HPRC-d46 graph and the haplotype-sampled HPRC-Sampled16 graph, then surjected onto CHM13. For Minimap2, these reads were mapped directly to CHM13 (see Section S5.3). Variant calling was done for both mappers using the ONT R104 model of DeepVariant for R10 reads, and the PACBIO model for HiFi reads mapped with Minimap2. For HiFi reads mapped with Giraffe, we used a custom model trained on reads mapped by Giraffe (see Section S5.4.1). Calling accuracy was evaluated using hap.py version 0.3.15 [37] against the Genome in a Bottle (GIAB) 4.2.1 truth set [38, 39]. We did not assess short read variant calling accuracy because this has been tested extensively [17, 18, 40] and has not changed with the new release of Giraffe.

Giraffe generally had slightly higher precision, recall, and F1 scores than Minimap2, with the haplotype sampling pipeline providing further improvements (Figure 5 A-D).

**Figure 5:**
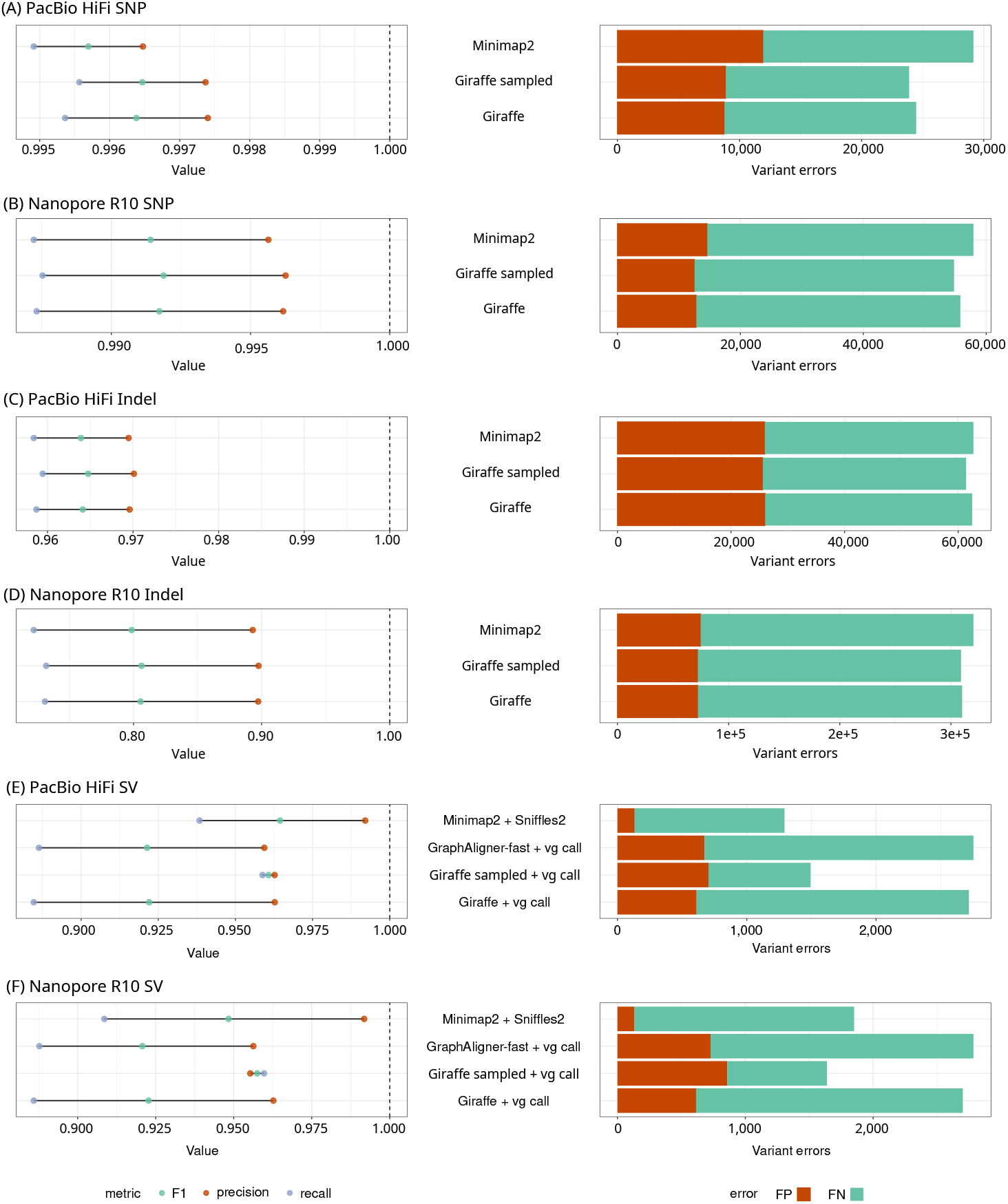
Variant calling and genotyping. SNPs (A and B) and indels (C and D) were called using DeepVariant with HiFi (A and C) and R10 (B and D) reads. SV genotyping was performed with Giraffe on the HPRC-d46 graph (Giraffe), Giraffe with a sampled graph with 16 haplotypes (Giraffe sampled), and GraphAligner’s fast mode on the HPRC-d46 graph, all with vg call. SV calling and genotyping was done using Minimap2 with Sniffles2. SV genotyping accuracy was compared for HiFi (E) and R10 (F) reads. Giraffe was used to map reads to the HPRC-d46 graph and to the HPRC-Sampled16 graph. Graphaligner’s fast mode was used to map reads to the HPRC-d46 graph. Minimap2 was used to map reads to CHM13.

**Figure 6:**
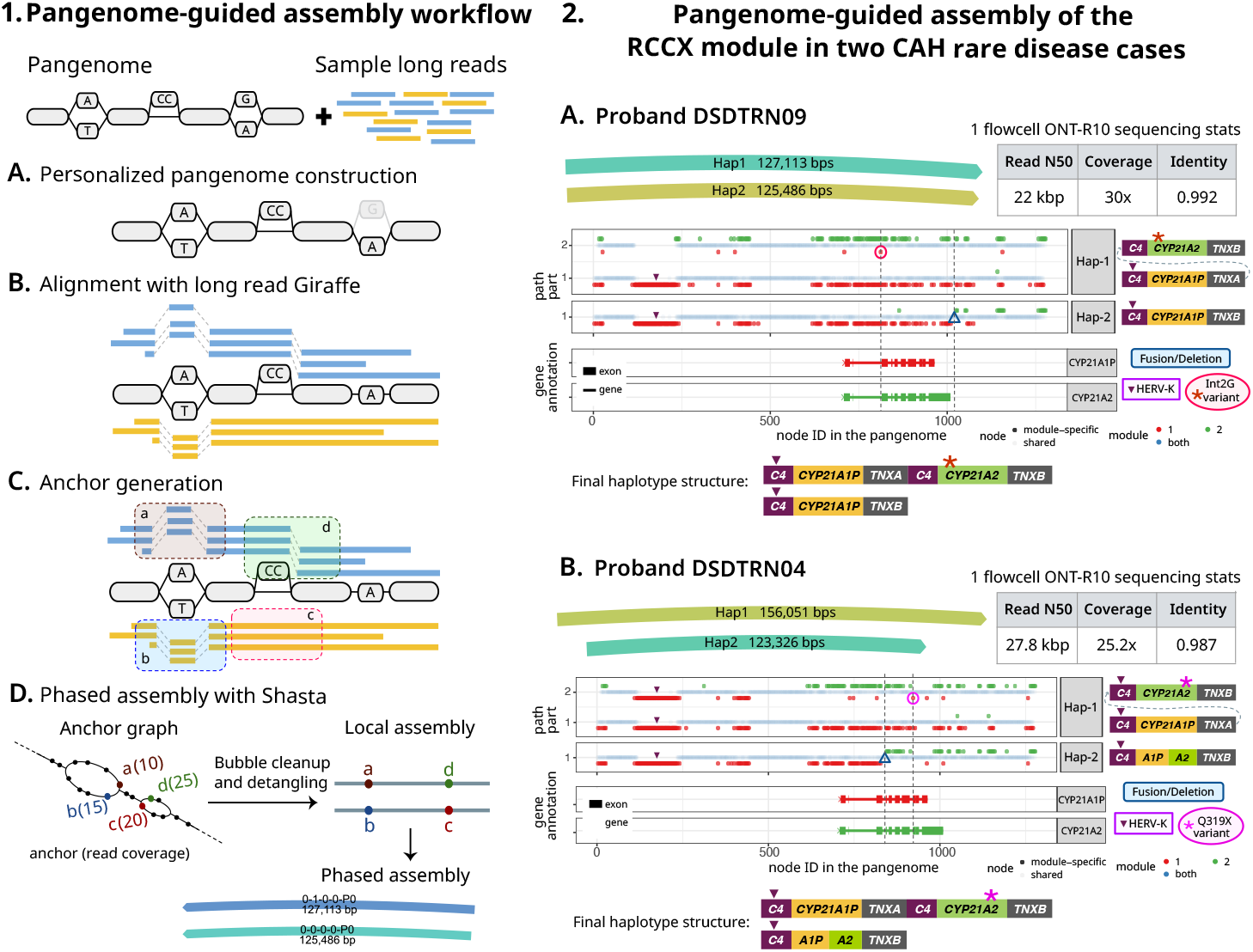
Pangenome-guided assembly workflow and application to rare disease cases.

### 3.5 Structural variant genotyping with vg call

We assessed the performance of Giraffe on structural variant genotyping with `vg call` using a pipeline based on Hickey et al. [41] (Section S5.5). `vg call` was run on Giraffe’s mappings to either the HPRC-d46 graph or the HPRC-Sampled16 graph. SV genotyping in each condition used the same graph that was used for mapping, except that for the HPRC-Sampled16 graph, a version was used with with both linear references included. We compared these two Giraffe conditions against two competitors: `vg call` on GraphAligner’s fast mode alignments to the HPRC-d46 graph, and the linear SV caller and genotyper Sniffles2 [42, 43] on Minimap2’s alignments to the CHM13 linear reference. Genotyping results were evaluated against the Genome in a Bottle Consortium’s T2T Q100 HG002 truth set [44] using Truvari [45] (Section S5.5).

Despite being restricted to only the variants present in the graph, the Giraffe-and-`vg call` pipeline with the haplotype-sampled graph performed comparably with Minimap2-and-Sniffles2, with the highest recall of any of the methods tested and a higher F1 score that Minimap2-and-Sniffles2 for R10 reads (Figure 5 E and F, Tables S13, S14).

### 3.6 Validation and application of pangenome-guided assembly for complex regions

We developed a prototype pangenome-guided assembler (PGA) (see Section 5.3) that leverages long read Giraffe alignments for targeted regional assembly. Briefly, Giraffe is used to map long reads to a pangenome. Sites of variation within the pangenome graph, termed bubbles or snarls, are used to determine haplotypic anchors in the reads, and then these anchors and their connections in the reads are used to create a phased genome assembly. Using the anchors derived from common variant sites, the assembly process is able to assemble a sample’s haplotype sequences, including private variations not present in the mapped-to pangenome.

We initially evaluated its performance using HG002 R10 reads (Section S6.1), across six selected loci: three structurally simple regions (CHR13-1Mb, CHR8-700kb, CHR2-600kb) and three complex, medically relevant regions with segmental duplications (RCCX, PDPK, GOLGA8).

For all six regions, PGA produced accurate phased assemblies (Figures S5S10). For comparison, we generated a complete Hifiasm assembly of the sample, aligned it to the HG002v1.0.1 reference with Minimap2, and retrieved the contigs overlapping the target region, trimming them to the aligned segments for evaluation. The evaluation showed comparable mismatch rates between both assemblies, even though PGA does not perform explicit read correction. In certain cases (Figures S6, S8), PGA produced better assembly contiguity than Hifiasm.

Following validation using the HG002 benchmark sample, we applied PGA to assemble the complex RCCX locus (Section S6.2) in two Congenital Adrenal Hyperplasia (CAH) cases, who had been analyzed in a previous study published by our group ([46, 47]), and compared the assemblies to those from a specialized method (Parakit) developed for analyzing this challenging locus [47]. With PGA, we were able to directly assemble the RCCX haplotypes for these two CAH probands (IDs DSDTRN09 and DSDTRN04 in Negi et al. [46]), and accurately detect the pathogenic compound heterozygous variants within these haplotype-resolved assemblies (Figure 6.2), recapitulating the earlier results made using Parakit and demonstrating accurate assembly of variants not found within the pangenome.

#### 1. PGA workflow overview

The inputs to the workflow are the reference variation graph and sample long reads. (A) Personalized pangenome construction using graph-unique k-mer classifications (faded node is not part of the personalized graph). (B) Reads are aligned to the personalized graph using long read Giraffe. (C) Read mappings at leaf snarls are used to generate anchors (potentially haplotype-informative k-mers enabling separation of reads). (D) The Shasta assembler constructs an anchor graph, with anchors as vertices and shared reads as edges. Its detangling algorithm traverses this graph to resolve haplotype paths and generate a phased assembly.

#### 2. Resolving RCCX haplotypes in two probands

For each proband, the figure displays PGA assemblies, sequencing statistics, the alignment profile of the phased haplotypes against an RCCX collapsed pangenome (using Parakit’s visualization), and the final RCCX haplotypes structure in sample. In the alignment profiles, when a haplotype traverses multiple RCCX modules (nodes: red = pseudogene-specific; green = gene-specific; blue = shared between SD modules) by looping through the pangenome, the different traversals are separated vertically and shown as multiple rows. The gene annotation track shows *CYP21A2* and *CYP21A1P* annotations. Vertical dotted lines highlight variant positions across panels. (A) Proband DSDTRN09: Hap-1 alignment through the pseudogenes followed by the genes reveals HERV-K deletion in *C4A* and a gene-converted pathogenic Int2G SNV (red circle). Hap-2 contains a fusion located beyond the *CYP21A2* gene in *TNXB* (blue triangle). (B) Proband DSDTRN04: Assembly alignment reveals a gene-converted pathogenic Q319X nonsense SNV (pink circle) in Hap-1 and a hybrid 5’-CYP21A1P/CYP21A2-3’ fusion in Hap-2.

## 4 Discussion

As the field of pangenomics continues to expand, there is a growing need for efficient tools for working with increasingly large and complex pangenome graphs. The latest version of Giraffe, part of our vg toolkit, is now capable of mapping both short and long reads to graphs of varying complexity—including to graphs constructed from more than 450 haplotypes from the HPRC’s second release.

For both short and long reads, Giraffe achieves similar read mapping, and overall slightly better variant calling accuracy, relative to existing best-of-breed linear and pangenome mappers, while being comparable in speed to the best linear mappers. Notably, Giraffe is over an order of magnitude faster than GraphAligner, the current state-of-the-art long read-to-pangenome mapper.

To demonstrate the potential of long-read-to-pangenome mapping, we developed a prototype pangenome guided assembly pipeline. With this pipeline, we were able to produce local, complete, phased assemblies of targeted regions of the genome, including those containing highly similar segmental duplications, that were more contiguous than the assemblies produced by Hifiasm and comparably accurate. Assembling the RCCX loci from two patients, we were able to accurately recover complex haplotypes that contain structural variations not present in the pangenome. This is in contrast to existing methods for structural variant calling against a pangenome, like the `vg call`-based methods evaluated here, which generally require the variation to be known in advance.

While further scaling and testing is required to convert this prototype assembler into a fully fledged tool, such a hybrid assembly process has great promise for future, efficient genome inference. It avoids the need for all-against-all read alignment (a step necessary during *de novo* assembly), and, by providing a strong prior for likely haplotypes, provides a framework that might facilitate more accurate assembly, particularly, we suspect, at lower sequencing coverages.

We demonstrate Giraffe’s performance on human graphs containing cycles and both large and small variants, but it is still untested on the more complex graphs constructed with the PanGenome Graph Builder (PGGB) [48] and on more genetically diverse species. PGGB’s reference-free approach to graph building provides an unbiased view of the homology within and among genomes. However, due to the complex topology of the PGGB graphs, we have not been able to construct indexes for Giraffe, and to our knowledge there are no tools capable of mapping reads to these graphs. There remains a need for indexes and algorithms that can handle the large cycles and deeply-nested variants present in PGGB graphs; we are actively working to extend Giraffe in this direction.

The construction of the graph being mapped to also has a profound effect on the downstream alignments and calling results. When graph alignments are projected back onto a linear reference, the base-level alignment of reads will often follow the alignment used to construct the graph. A pathological “spelling” of the graph’s alignment (for example, one providing multiple paths that spell the same sequence) may negatively affect downstream results, even if the read was mapped to the correct location and represents the correct sequence.

We found ourselves obliged to work around this effect (see Section S4.2). We are actively pursuing new graph construction techniques that we think will allow for building graphs that produce graph alignments that in turn produce more useful linear alignments. We also suspect that there may be an opportunity here to revive the field of indel realignment, seemingly last popular in the era of GATK 3. We hypothesize that an indel realigner with long read support, ideally working in pangenome space, would improve point variant calling accuracy from pangenome alignments. Unfortunately, we know of no suitable existing tool.

There is still work to be done to further improve the Giraffe algorithm. We currently use the original Giraffe algorithm for mapping paired-end short reads, but we expect results to improve when adding pairing support to the long read algorithm. Furthermore, while the improvements in variant calling demonstrated are currently modest, we believe that there is considerable room for further improvement. One future idea is to score recombinations, and so penalize read mappings that are haplotype inconsistent, as is often case when mapping reads to paralogs [31]. Such recombination awareness also opens the possibility to assign reads to local haplotypes and use these local haplotypes more directly in downstream applications.

## 5 Online Methods

### 5.1 Variation graphs and indexes

Giraffe is implemented as part of the vg toolkit, which uses a **variation graph** model for representing pangenomes [7]. A variation graph is a bidirected graph in which **nodes** are labeled with nucleotide sequences, and **edges** represent possible adjacencies between these sequences. Nodes in a variation graph have two sides, which are arbitrarily designated as **left** and **right**. Edges connect pairs of node sides. Valid walks through the graph must enter and exit opposite sides of a node, so each node visit along the walk has an orientation. A forward (i.e. left to right) traversal of a node represents the forward sequence of the nucleotides, whereas a backwards traversal represents its reverse complement. Walks through the graph can therefore be used to represent longer sequences, found by concatenating the sequences of the nodes. We call walks used for this purpose **paths**, and they are usually stored along with the variation graph that they are inside of.

Paths in a variation graph usually correspond to a small number of reference sequences. In addition to the reference sequences, Giraffe also uses a larger panel of haplotype paths, stored in a **Graph Burrows-Wheeler Transform** or **GBWT** index [49]. Originally, GBWT indexes were stored separately from their graphs, but the **GBZ** format [50] was developed to store them together, by storing the GBWT and a “GBWT graph” of node sequences in the same file, and storing non-haplotype paths along with the haplotypes in the GBWT. Because we found GBZ graphs so easy to work with, Giraffe now always uses the GBZ format. One subtlety is that a GBZ uses the paths in the GBWT to implicitly define the edges of the graph. Thus, a GBZ graph represents the subgraph of some original sequence graph whose edges are supported by the haplotypes and reference paths in the pangenome.

Giraffe uses a distance index [51] to find minimum distances between positions in the graph. The distance index is based on decomposing the graph into **snarls** [52], which are sites of variation in the graph where paths can diverge to represent different alleles.

Formally, a snarl is a subgraph defined by two boundary node sides that are **separable** and **minimal**. Two nodes sides are **separable** if cutting their nodes into their component node sides makes the subgraph between them unreachable from the rest of the graph. The two boundary node sides are **minimal** if no other node side within the subgraph is separable with either of the boundaries. Snarls often occur consecutively, with boundaries on opposite sides of shared nodes between them.

A maximal length run of one or more nodes, with snarls between each node and the next, is called a **chain**. (These chains of snarls are not to be confused with the chains of alignment seeds used later.) The simplest possible chain is just a single node, so every node exists in some chain. More complex chains can be nested within snarls to represent nested variation, such as a series of SNPs that occur within an insertion. The decomposition of the graph into nested snarls and chains is described by a snarl tree (Figure 1 B). Although formally the decomposition is not unique, we usually work with one decomposition at a time.

### 5.2 Giraffe algorithm

The long read Giraffe algorithm follows the common seed-chain-extend strategy [11] for mapping algorithms (Figure 1). As in the original short read algorithm, long read Giraffe uses a minimizer index over the sequences of haplotypes in the GBWT to find seed alignments (Figure 1 C). For long reads, we use weighted minimizers that prioritize those with fewer occurrences in the graph, and add a filter to skip minimizers with too many occurrences (Section S1).

The seeds are then chained together (Figure 1 D-E, Section S3). A **chain** of seeds (not to be confused with the chains of snarls used previously) is an ordered set of seeds that are co-linear in the read and the reference. In a graph context, this means that each seed must be reachable in the graph from the previous seed. Giraffe tries to find optimal chains that maximize the coverage in the read and minimize the expected cost of the **gap** (the difference between the distance between the seeds in the read and the distance between seeds in the graph) that will be taken between consecutive seeds. We use a gap cost based on that of Minimap2 [10].

To efficiently calculate reachability and distance in a graph, we use a novel data structure called a **zip code tree** that represents distances among a set of seeds (Figure 1 D). The zip code tree approximates a placement of seeds in an unrolled, acyclic view of the reference graph, providing a partial ordering of seeds that can be used to find distances for chaining (Figure S1, Section S2). Using distances calculated with the zip code tree, we are able to find reasonable chains in regions of the graph with complex topologies such as duplications (Figure S2) and inversions (Figure S3). The zip code tree is computed using small data structures called **zip codes**, that store, for each seed, information about the position’s placement on the snarl tree and associated distances (Section S2.3). Because of the cache-efficiency of these lightweight data structures, it is fast to sort and calculate distances among seeds to construct the zip code tree. A zip code tree is constructed for all seeds in a single read. Chaining is done using two passes of a dynamic programming algorithm (Section S3.1).

Finally, the chain (Figure 1 E) is extended into an alignment between the read and the graph. Base-level alignment is done between consecutive seeds in the chain and out from the first and last seeds (Figure 1 F, Section S4). We use a fast wave-front algorithm [53, 54, 55] for aligning substrings of the read to the haplotype sequences in the graph, both for between-seed alignment and anchored tail alignment. If this algorithm fails to find an alignment or is predicted to use too much memory (or for any alignment problem over 233 bases), we align to the graph itself, using a banded global aligner for aligning between seeds and a Single Instruction Multiple Data (SIMD)-accelerated X-drop aligner (called Dozeu) for aligning tails. These algorithms limit the size of the dynamic programming band to constrain the amount of time and memory used.

### 2.3 Pangenome-guided assembly overview

Using Giraffe’s ability to align long reads to pangenomes, we can do haplotype-resolved pangenome-guided assembly of long reads. Here we focus on local assembly.

Our method assembles a specified region of interest (ROI) using a personalizedpangenome-guided assembly workflow, depicted in Figure 6.1. The workflow takes as input a variation graph and the sample’s long reads. For this study, we used the HPRC-clipped pangenome graph.

First, the variation graph is personalized based on the input reads [18] by sampling 16 haplotypes (without using `--diploid-sampling,` which would further downsample to two). Input reads are then mapped to this personalized graph using long read Giraffe. For a target ROI, all reads mapping to the region, along with the subgraph and its snarl distance index, are extracted. These serve as input for the subsequent anchor generation step.

**Anchors** are defined as potentially haplotype-informative k-mers, and enable partitioning of reads by haplotype. They are generated from unique paths within **leaf snarls** (snarls that contain no other nested snarls). Anchors of a given snarl share common boundary nodes but differ at internal sentinel node(s) that represent allelic variation. An anchor is considered to be supported by a read only if the corresponding read subsequence aligns as an exact match to the anchor path. To mitigate phasing errors caused by noisy long read alignments, we perform a filtering step to remove anchors in **unreliable snarls** that exhibit inconsistent phasing relative to neighboring snarls. Finally, the complete set of read-supported anchors from the reliable snarls is passed to the Shasta assembler, which performs bubble cleanup and detangling to generate a phased assembly of the ROI.

## 6 Acknowledgments

## 6.2 Author contributions

Project design: B.P. Giraffe algorithm design and implementation: X.C., A.M.N., J.M.E., J.S., S.H., B.P. Pangenome-guided assembly: S.Ne., F.A., S.Na., K.K. Graph production: G.H. Structural variant analysis: J.M., X.C., A.M.N. Short variant analysis: A.C., K.S., P.-C.C., A.M.N., X.C. vg code contributions: A.M.N., X.C., J.M.E., J.S., J.M., G.H., F.O. Sample collection from DSD-TRN biobank: E.D.

## 6.4 Data and materials availability

An archived copy of the code, graphs, DeepVariant models, and simulated reads (excluding R10 reads) used in this paper are available on Zenodo [56]. All data on Zenodo is also available at https://cgl.gi.ucsc.edu/data/lr-giraffe/, along with the R10 simulated reads and HG002 real read sets used for mapping evaluations.

These real read sets used for mapping and calling valuations are available on Zenodo, at https://cgl.gi.ucsc.edu/data/lr-giraffe/, and are publicly available at https://s3-us-west-2.amazonaws.com/human-pangenomics/index.html?prefix=T2T/HG002/assemblies/polishing/HG002/v1.0/mapping/hifi_revio_pbmay24/ (HiFi reads), s3://ont-open-data/giab_2025.01/ (R10 reads), gs://brain-genomics-public/research/sequencing/fastq/novaseq/wgs_pcr_free/40x/ (Illumina reads), and https://s3-us-west-2.amazonaws.com/human-pangenomics/index.html?prefix=T2T/scratch/HG002/sequencing/element/trio/HG002/ins500_600/ (Element reads). The read sets used for training DeepVariant are also publicly available; for details see Section S5.4.1.

The HG002 R10 reads used for the pangenome-guided assembly pipeline are publicly available from GIAB here https://epi2me.nanoporetech.com/giab-2025.01/. The CAH samples are available from the authors upon request (see Section S6.1).

The HPRC-d46 graph and HPRC-Minigraph graphs used for mapping and calling evaluations are available on Zenodo [56], at https://cgl.gi.ucsc.edu/data/lr-giraffe/, and at https://s3-us-west-2.amazonaws.com/human-pangenomics/index.html?prefix=pangenomes/scratch/2025_02_28_minigraph_cactus/. The personalized haplotype sampled graphs are not publicly available, but they can be recreated using HG002 read sets and the base HPRC-clipped graph, which is available on Zenodo [56] and at https://cgl.gi.ucsc.edu/data/lr-giraffe/.

The Snakemake workflow used for read mapping, alignment analyses, and variant calling is available at https://github.com/vgteam/long-read-giraffe-experiments/tree/main. The regional PGA Snakemake workflow used for HG002 testing is available at https://github.com/shlokanegi/pga_workflow/tree/lrg2025-paper. The latest version of the vg toolkit is released at https://github.com/vgteam/vg. Copies of each of these GitHub repositories are available on Zenodo [56].

