## Supplementary Materials for "Rapid, accurate long- and short-read mapping to large pangenome graphs with vg Giraffe"

#### Part S
### Supplementary Material

#### S1 Seeding

Seeding is done using a minimizer index of the haplotypes in the GBWT. We used weighted minimizers with a k-value of 31 and a window length of 50 for aligning long reads. Minimizer weighting prioritizes minimizers with fewer hits in the reference similar to Jain et al. [57]. For short reads, we use unweighted minimizers with a k-value of 29 and window length of 11.

Because long reads produce a large number of minimizers, each of which may have many occurrences in the graph, we filter the seeds to pare them down to a workable number. As in the original Giraffe algorithm, we have the option to use a *hit-cap*, *hard-hit-cap*, and *score-fraction* to filter seeds based on the number of occurrences (hits) they have in the graph. If a minimizer has fewer than *hit-cap* hits, then it is always included. If it has more than *hard-hit-cap* hits, then it is always discarded. For seeds whose hit count is between the two caps, we go through the minimizers in increasing order of hit count and accept minimizers until the sum or their scores is *score-fraction* of the total score of all minimizers found. As in the original Giraffe algorithm, the score of a minimizer is defined as $max(1 + ln(C_h - ln(n)), 1)$, where $C_h$ is the *hard-hit-cap* and and $n$ is the number of hits of the minimizer [17]. Additionally, the total number of minimizers used is capped at the maximum of *max-min* and the read length divided by *num-bp-per-min*. For long reads, the *hit-cap* and *score-fraction* are ignored in favor of a downsampling approach. In this approach, we pass a sliding window across the read and, for each window, keep only the seed with the fewest number of hits in the graph. The length of the sliding window is the minimum of the *downsample-window-length* and the length of the read divided by the *downsample-window-count*

#### S2 Zip Code Trees

##### S2.1 Overview

The problem of co-linear chaining of seeds in a graph requires a concept of reachability and distance, both of which are non-trivial problems in a graph context. In previous versions of Giraffe, we used a snarl decomposition-based distance index [51] to calculate minimum distances in the graph. While theoretically efficient, in practice calculating distances from the index could be very slow due to cache misses when querying parts of the index that were located far apart in memory. To address this issue, we designed a new data structure called a **zip code** to store information from the distance index for a node in the graph. Similar to how a real-life zip code is used to specify the location of a mailing address from the larger geographical area to the more specific, our snarl-based zip codes specify the location of a node on the snarl tree, starting from the largest root-level structure going down the tree to the leaves. At each level of the snarl tree, the zip codes store an identifier for the snarl or chain and enough information to calculate distances. By comparing the zip codes of two nodes, we can quickly determine their common ancestors and find the minimum distance between them. Due to their small size, calculating distances with the zip codes has better memory co-locality than doing so with the distance index.

We wanted to leverage the fast distance comparisons of the zip codes to support the distance and reachability queries needed for chaining. When chaining together seeds, we are generally interested in finding relatively small distances between seeds that are near each other in the graph, and we don't care about very long distances between seeds that are far from each other as these seeds are unlikely to be in the same chain. We therefore wanted a data structure that could quickly determine the distance between nearby seeds. We call this data structure a **zip code tree**.

The zip code tree can be thought of as a subgraph of the original variation graph containing only the nodes that contain seeds, and a minimal number of additional nodes and edges to connect the seeds. These additional nodes represent the boundaries of snarls and chains in the snarl tree, based the same intuition underlying the distance index: that minimum-length paths between nodes must include the minimum-length paths through the bounds of containing snarls and chains [51]. Edges in the zip code tree are labeled with minimum distances from the original variation graph. Minimum distances between seeds can therefore be found in a shortest-path traversal of the zip code tree. Although this would be slow for very long distances, it is an efficient way to find the relatively short distances that we are interested in for chaining.

A zip code tree is constructed for each read individually. Zip code tree construction is done by sorting the zip codes of seeds in a radix-like sort of their positions along the snarl tree, then adding snarl and chain boundaries and distances between nodes. This process is fast due to the low computational cost of comparing zip codes. Separate trees are found for seeds that are unreachable or that are farther than a given distance limit from other trees. Similar to a cluster of seeds, each disconnected zip code tree represents a distinct region of the graph that is a potential placement of the read. Distances between seeds are found for the chaining step by iterating through the zip code tree (Section S2.5). Chaining is done separately for each disconnected zip code tree.

#### S2.2 Zip code tree structure

Zip code trees are organized using the snarl decomposition of the graph(Figure S1 A). Each node in the zip code tree represents the position of a seed in the graph or the boundary of a snarl or chain containing a seed (Figure S1 B). Each seed is associated with a zip code. Edges occur between pairs of zip code tree nodes that are parents/children or siblings in the snarl tree; edges are labeled with graph minimum distances between the structures they represent. Thus, the zip code tree can be viewed as representing the same layout of the original graph, including only the seed positions and the boundaries of their containing snarls and chains. The distances between seeds can be found by walking through the zip code tree and taking the sum of the edge weights (Figure 1 D).

(Note that the structure of "nodes" described here is *not* a tree. The "zip code tree" gets its name because the initial ordering of the nodes, found in a radix-like sort of the zip codes, follows a pre-and-post-order traversal of the snarl tree structure. In this way, the zip codes are organized into a suffix tree that is flattened with additional edges added. The actual edges in the zip code tree do not form a tree.)

Zip code trees are stored as a single vector of nodes and edges. Nodes and edges therefore have a fixed orientation in the zip code tree, which may not match that of the original graph. Edges always connect the local right side of a node to the local left side of the downstream node. This imposes the restriction that zip code trees are directed acyclic graphs (DAGs) and cannot represent the full connectivity of arbitrary variation graphs. When constructing zip code trees for graphs that are not already DAGs, we use the order and orientation of the seeds in the read to transform the graph into a DAG that best represents a likely path of the read (Figures S2 and S3).

The final form of a zip code tree is a linearized sequence of **symbols**, corresponding to its nodes (`SEED`, `CHAIN_START`,`CHAIN_END`, `SNARL_START`, or `SNARL_END`), edge distances between them (`EDGE`), or a count of children in a snarl (`NODE_COUNT`) (Figure S1 B). This sequence can then be used to list the possible predecessors of each seed (see S2.5).

##### S2.2.1 Chains

Chains in the zip code tree are comprised of their child snarls and seeds that occur on node children of the chain (Figure S1 C). All seeds are contained within a chain, even if it is the only seed in the chain. For chains, there is an edge from the start bound of the chain to the first seed in the chain, an edge between each consecutive seed or child snarl, and an edge from the last child of the chain to the end bound of the chain. Edges involving snarls terminate at a bound of the snarl.

In the vector representation of a chain in the zip code tree, chains start with a `CHAIN_START` symbol and end with a `CHAIN_END` symbol. Inside the chain, there are child `SEED` symbols and snarls (which are similarly bounded by `SNARL_START` and `SNARL_END` symbols), and `EDGE` symbols between each consecutive child snarl or seed.

##### S2.2.2 Snarls

The representation of a snarl in a zip code tree depends on its topology. We classify snarls as **regular**, **irregular**, or **cyclic**. A snarl is regular if all of its children have only two edges: an edge from one side of the node to the start or end bound, and an edge from the opposite side of the node to the other bound. The only other edge allowed in a regular snarl is between the two bounds. A non-regular snarl can be classified as cyclic if there exists a path from any node (including boundary nodes) back to itself on either side. Note that this definition allows for "cyclic" snarls that do not actually have any cyclic walks (returning to the same orientation of the same node). All non-regular, non-cyclic snarls are classified as irregular. In Figure S1 A, the orange snarl is a regular snarl and the green snarl is an irregular snarl. The two snarls in Figures S2 and S3 are both classified as cyclic snarls.

For regular and irregular snarls (but not cyclic snarls) in the zip code tree, each child can only be traversed in one direction in a start-to-end traversal of the snarl. That is, there exists a path from the start boundary of the snarl to one side of the child and a path from the end boundary of the node to the other side of the child, but no path from the end boundary node to the first side of the child or from the start boundary node to the second side of the snarl. We will arbitrarily refer to the side of the child reachable with the start boundary node as the **left** side and the side of the child reachable with the end boundary as the **right** side. This may be different from the orientation assumed in the distance index. Additionally, since these snarls are DAGs by definition, there exists a topological ordering of snarl children. For regular and irregular snarls, the zip code tree contains an edge from the snarl start bound to the left side of every child, and an edge from the snarl end bound to the right side of every child. For irregular snarls, there is also an edge from the right side of every child to the left side of every other child succeeding it in the topological order (Figure S1 D).

The vector representation of a snarl starts with a `SNARL_START` and contains child chains and edges between the chains and bounds (Figure S1 D). Note that a chain is contained entirely within a snarl, so edges terminating at chains are really terminating at the `CHAIN_START` or `CHAIN_END`. The next symbol in the vector is an `EDGE`, and is followed by the first chain in the snarl (starting with the `CHAIN_START` and ending with the `CHAIN_END`). This edge represents the distance from the snarl start bound to the first chain. If there is a second chain, then the next symbol would be an `EDGE` representing the distance from the first chain to the second chain, followed by another `EDGE` representing the distance from the start bound to the second chain, followed by the second chain. In this way, each chain is preceded by an `EDGE` representing the distance to every node symbol before it in the snarl, in reverse order that they appear in. This holds true for the `SNARL_END` as well; it is preceded by an edge for each chain in the snarl and for the `SNARL_START`. Note that an `EDGE` may contain a distance value of infinity, indicating that there is no valid walk between the nodes connected by the edge. The second-to-last symbol in the snarl, after the `EDGE` between the two bounds and before the `SNARL_END`, is a `NODE_COUNT` that represents the number of chains in the snarl.

##### S2.2.3 Cyclic snarls

Cyclic snarls are not DAGs and may not have a topological order or a single possible orientation of children. To account for how paths through a cyclic snarl may traverse the snarl's children multiple times and in multiple directions, we may represent the same chain multiple times in the zip code tree. Since a zip code tree is required to be acyclic, we cannot visit the same representation of a chain more than once in a traversal of the zip code tree, so we need multiple copies. We want to represent each chain enough times to represent all paths through the snarl that are possible in an alignment between the read and graph. We use the seeds' positions in the read to estimate the required copy number and orientation of each child chain.

Seeds in the snarl are split up into **runs** of seeds that are on the same child chain in the same orientation on the read and within a given distance limit of each other in both the read and chain. Each run of seeds is considered to be a separate child of the snarl in the zip code tree, even though there may be multiple runs on the same child chain in the underlying graph's snarl tree. Rather than sorting the runs by topological order, as we do for non-cyclic snarls, runs are sorted according to the seeds' positions in the read. This order is treated as a topological order for an irregular snarl, and edges are added accordingly (Figures S2 and S3).

Some child chains of cyclic snarls may be traversed in either direction in a start-to-end traversal of the snarl, but each run can only be written out in one orientation at a time. The orientations used for runs in the zip code tree are determined by the order of the seeds in the read (Figure S3). First, we find the Spearman correlation between the coordinates of the positions in the read and in the snarl's parent chain, to determine if the read should traverse the snarl forwards or backwards. Next, we find the same correlation for each of the runs. If there is a strong enough correlation, we orient the run so that the read can traverse the run and the snarl in the same direction. If there isn't a strong correlation, we keep a copy of the run in each direction so it can be traversed in either orientation.

Using this method, there may be multiple instances of a child of a snarl represented as separate runs of seeds, and there may be up to two copies of each child, one in each orientation. Because cyclic snarls may be nested inside each other, duplicating every child in every nested cyclic snarl could result in an exponential blowup of copies of the child. To prevent this, we only allow one copy of a child to be duplicated again in a more deeply nested snarl. The other copy of the child is treated as a chain in a non-cyclic snarl and is not permitted to duplicate itself. This strategy tends to split the chains into more runs than are necessary to represent cyclic paths, but it tries to ensure that all probable paths through the snarl are found.

#### S2.3 Zip codes

When constructing zip code trees, we rely on the zip codes associated with each seed to quickly find the placement of the seed in the snarl tree, and to calculate distances among seeds and snarl tree boundaries. The original Giraffe algorithm calculated distances using a minimum distance index based on the structure of the snarl decomposition [52]. Walking up the snarl tree, the minimum distance

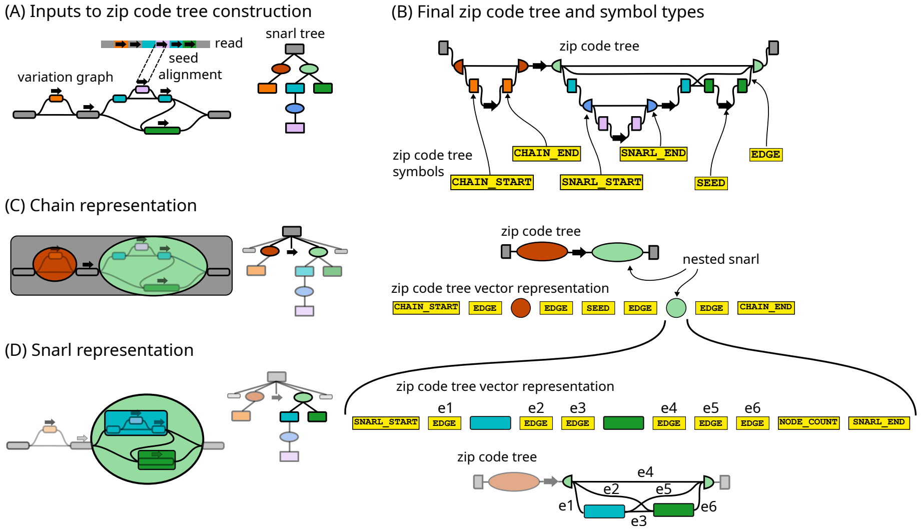


Figure S1: **Zip code trees**. (A) A sequencing read, variation graph, and corresponding snarl tree from Figure 1. Arrows on the variation graph and read represent seed alignments. Rectangular nodes in the snarl tree represent chains and round nodes represent snarls. Chains are colored by the nodes in the chain they represent on the graph. For simplicity, the snarl tree does not represent nodes, which are children of chains. (B) The zip code tree for seeds on the graph in (A) and a subset of the corresponding symbols that are stored in the vector representation of the zip code tree. Nodes in the zip code tree represent either seeds (arrows) or the start or end bounds of snarls and chains. The bounds are colored and shaped like the snarl tree nodes they represent. (C) The representation of the top-level chain (grey) in the variation graph, snarl tree, and zip code tree. The chain has five children: two snarls and three nodes, one of which has a seed on it. The zip code tree representation of the chain contains the start and end of the chain, all snarls that contain seeds, and all seeds on node children of the chain. The zip code tre vector representation shows the order of symbols actually stored in the vector. (D) The representation of a snarl in the graph, snarl tree, and zip code tree. The light green snarl has two children: the blue chain and the green chain. In the zip code tree, the two chains are sorted according to a topological order of the snarl subgraph. Edges occur from the right side of each child of the snarl to the left side of every child later in the topological order. In the full zip code tree, the vector representation shown here would replace the green circle in the vector representation in (C).

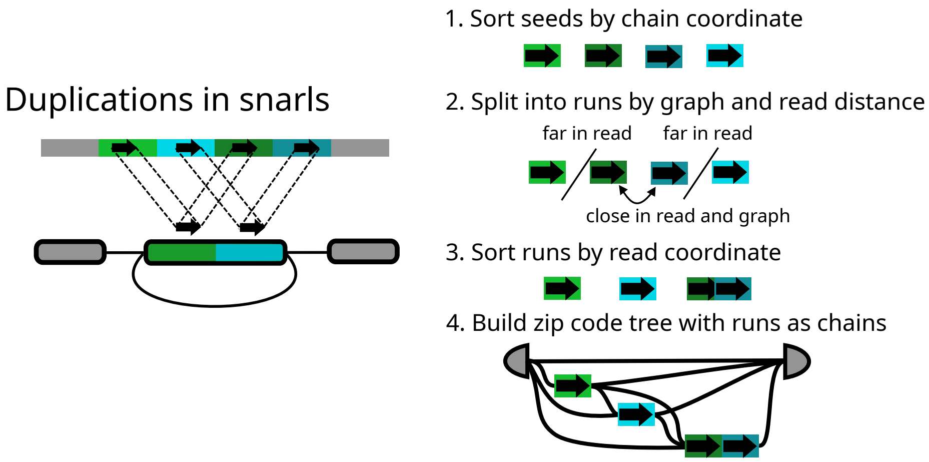


Figure S2: **Duplications in zip code trees**. Seeds in the chain are split into runs of seeds based on the distances between the seeds, each of which represents a separate potential traversal of the duplicated chain. Each run becomes a separate child of the snarl.

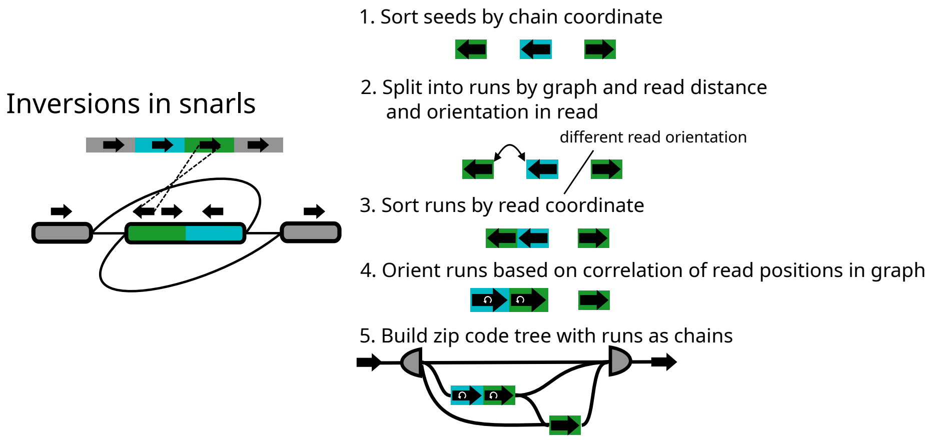


Figure S3: **Inversions in zip code trees**. Seeds in the chain are split into runs based on the distances between them and their orientations in the read. The orientation of a run is determined by finding the expected orientation of the read through the snarl, based on context in the snarl's parent chain. Seeds that are oriented backwards in the zip code tree relative to the forward orientation of the node they are on are indicated with a white arrow.

from each position to the bounds of each ancestor snarl or chain is found. At each common ancestor of the two positions, the distance between the children (or child) of the common ancestor containing the two positions is found and added to the relevant distances to bounds of the child to find the minimum distance between the positions in this common ancestor. This step must be done for every common ancestor, not just the lowest common ancestor, because the minimum distance path may take edges outside of the lowest common ancestor.

Zip codes store a small amount of information about a graph node, and allow calculating minimum distances between any two nodes in the graph, given their zip codes and (in some cases) the distance index information for their common ancestors in the snarl tree. Unlike approaches such as hub labeling [58], zip codes leverage the fact that pangenome graphs are likely to have a "good" snarl decomposition, with many long chains, and the zip codes are easy to compute from that decomposition. The basic structure of a zip code is a vector of **codes**, with one code per snarl tree node (snarl, chain, or graph node itself) that is an ancestor of the graph node, starting from the root structure and going down to the node. Each code contains an identifier for the snarl tree node and distance information to either find distances directly or to query the distance index. Identifiers are only unique for structures with the same ancestors, i.e., two chains that share a parent snarl will always have different identifiers, but two chains in different snarls may have the same identifier. In this way, the series of identifiers of a zip code, starting from the root and proceeding down to the node, is sufficient for uniquely identifying the node and for finding the common ancestors of two nodes.

#### S2.4 Zip code tree construction parameters

Zip code trees are only guaranteed to store distances up to a given distance limit. The **distance limit** is the length of the read times the *zipcode-tree-scale*. If all seeds in a chain or a segment of a chain are farther than the distance limit away from anything else in the chain or containing snarl, then the chain or chain segment is separated out into a separate zip code tree. This helps reduce the size of individual zip code trees, and thus individual chaining problems.

We filter zip code trees based on their score and coverage. The **score** of a zip code tree is the sum of the scores of the minimizers that it contains. Similarly, the **coverage** of a zip code tree is the fraction of the read that is covered by the minimizers in the tree. A zip code tree is filtered out if its score is more than *zipcode-tree-score-threshold* worse than the best-scoring zip code tree, or if its coverage is more than *zipcode-tree-coverage-threshold* less than the zip code tree with the best coverage. We keep at least *min-to-fragment* and at most *max-to-fragment* zip code trees.

#### S2.5 Zip code tree iteration

The zip code tree is used to find **transitions** between seeds by identifying pairs of "source" and "destination" seeds connected by sufficiently short paths in the graph.

We iterate through the zip code tree vector from left to right. When we encounter a seed in the zip code tree vector, we find the corresponding "anchor" in the chaining problem (see Section S3). If there is no such anchor, we skip the seed.

We then consider that seed to be the **destination**, and iterate from there right to left looking for possible **source** seeds and their associated graph distance estimates (**graph distances**), stopping when we know that no more possible sources with graph distances within the **graph lookback limit** can be found. The graph lookback limit is computed as the maximum of a flat value and a per-base value times the length of the read being aligned, with these parameters set separately for each round of chaining (see Section S3.2).

##### S2.5.1 Pushdown automaton iterator

To actually enumerate the potential source seeds for a destination seed in the zip code tree vector, an iterator widget is implemented in the form of a **pushdown automaton**, which is a finite state machine (itself a classic computer science concept that consumes a sequence of input symbols), extended with a stack. This iterator maintains a state, to track what it is currently doing, and a stack, to track several running distances and return to older ones as iteration through chains within a snarl finishes. It is responsible for yielding possible source seeds and their graph distances from the destination seed.

The automaton uses a notion of adding distances where there is an infinite-distance value (represented as an all-`1` binary value), and adding anything to an infinite distance produces an infinite distance.

The basic idea is that the automaton tracks what kind of structure it is in (chain or snarl) and what it is doing with it ("stacking" up distances, "scanning" for seeds to yield, or "skipping" over seeds known to be too far from the destination) using its state, and jumps between states depending on its current state, the symbols it encounters in the zip code tree (reading right to left), and the contents of its stack. The stack will mostly hold various forms of **running distance**, which is the minimum distance between wherever the iterator is in a snarl or chain and the destination seed. We will now review the specific states and transitions.

`START` **state** The automaton starts in `START`, with an empty stack, pointed at a `SEED` as the next symbol, corresponding to the seed to traverse left from. On processing the symbol, the automaton sets up the stack by pushing a zero running distance, and proceeds to `SCAN_CHAIN`.

`SCAN_CHAIN` **state** In this state, the automaton is scanning a chain leftward towards its start, visiting seeds or contained snarls. The stack holds the running distance along the chain as its top element. Below that, for each level of enclosing snarl that was traversed into, the stack will contain the running distance to use when traversing each chain in that snarl that's earlier in the zip code tree than the current position, and the end-to-start distance through that snarl.

When an EDGE is encountered, it represents a distance between two adjacent things along the chain being scanned. Its stored distance is added into the current running distance at the top of the stack. If this exceeds the distance limit being used for the search, the rest of the chain needs to be skipped. The automaton checks the stack depth to determine if the parent snarl of the current chain is being traversed. If so, it pushes a zero to the stack, representing the depth of nested snarls currently being skipped over, and goes to SKIP_SNARL. If not, the automaton halts, since nothing to the left along the chain or in the enclosing snarl can be within the distance limit.

When a SEED is encountered, it is emitted as a potential predecessor, using the running distance from the top of the stack.

When a SNARL_END is encountered, the automaton goes to the STACK_SNARL state, to begin collecting the distances required to traverse the child chains inside the snarl.

When a CHAIN_START is encountered, the chain has been scanned completely. The automaton again checks the stack depth to determine if the parent snarl of the current chain is being traversed. If so, the automaton pops and discards the running distance along the current chain, and goes to SCAN_SNARL to continue using the running distances to things in that snarl that are currently on the stack. Otherwise, the running distance is kept, because the automaton started its traversal within the current chain, and the automaton goes to STACK_SNARL to use the stored distances to other things in the snarl, and the start of the snarl, that appear before the chain in the zip code tree.

STACK_SNARL **state** In this state, the automaton is stacking up the stored distances to things within a snarl. The stack contains the running distance, either along the parent chain of the snarl or along the child chain from which the snarl is being entered. This is at the top of the stack so it is available to add to distances read from the zip code tree. Below that are all the running distances within the snarl that have already been stacked. Below them, the stack contains similar running distance sets for each level of enclosing parent snarl that was traversed into.

When an EDGE is encountered, it represents the distance from the snarl end, or from the chain being read out of, to some earlier thing in the snarl, or to the snarl start. The automaton duplicates the running distance at the top of the stack, adds the symbol's stored distance to the top element of the stack, and then swaps the top 2 stack elements, leaving a new running distance to the edge's destination on the stack, below the unchanged top running distance.

When a CHAIN_END symbol is encountered, the automaton knows that the complete table of distances within the snarl has been incorporated into the stack. It pops and discards the top running distance, and then checks the stack depth. If it finds the stack to be empty, it concludes that it has read out the left end of a top-level chain and found a different, completely disconnected top level chain, and halts. (This allows one zip code tree data structure to describe multiple disconnected components, or multiple orientations of the same connected component that should not interact.) If the stack is not empty, it contains a running distance to be used when traversing this newly encountered chain. If that running distance exceeds the distance limit, the automaton pushes a zero, representing the depth of nested snarls currently being skipped over, to the stack, and goes to `SKIP_SNARL`. Otherwise, it leaves that running distance at the top of the stack and goes to `SCAN_CHAIN`.

When a `START_SNARL` symbol is encountered, the automaton checks to see if the running distance is the only thing on the stack, or if it has available a stacked distance that corresponds to the snarl start. If only the running distance is on the stack, it concludes that it has read to the start of a top-level snarl, containing only disconnected top-level chains, and halts. If a distance on the stack for the snarl start is available, it pops the running distance off the top of the stack, leaving the distance for the snarl start at the top, and goes to `SCAN_CHAIN` to scan along the snarl's parent chain.

`SCAN_SNARL` **state** In this state, the automaton is consuming the distances from the stack that were stacked up while in `STACK_SNARL`. The stack contains distances to use as running distances when scanning each chain to the left of where the automaton is in the snarl, and the distance for the snarl start (to be used to continue along the snarl's parent chain), in the order that the automaton will encounter the corresponding objects. Below them, the stack contains similar distances for each level of enclosing parent snarl that the automaton read into.

When a `CHAIN_END` is encountered, the automaton checks the distance for it at the top of the stack, to see if it is over the distance limit. If so, the automaton pushes a zero, representing the number of nested levels of snarls currently being skipped, to the stack, and goes to `SKIP_CHAIN`. Otherwise, it keeps that running distance for the chain at the top of the stack and goes to `SCAN_CHAIN`.

When a `SNARL_START` is encountered, the automaton already has the appropriate distance on the top of the stack to become the running distance along the snarl's parent chain, so it leaves it there and goes to `SCAN_CHAIN` to scan that parent chain.

When an `EDGE` is encountered, it is ignored; it represents distance information that is only relevant for coming out of other child chains within the snarl.

When a `NODE_COUNT` symbol is encountered, it is also ignored. Snarls in the zip code tree can contain node count information, but the pushdown automaton iterator does not use it.

`SKIP_CHAIN` **state** This state is used when the automaton has determined that everything to the left of a certain point in a chain will have a distance over the distance limit, but that more results withing the distance limit might be available from sibling chains in an enclosing parent snarl. The top of the stack holds the number of levels of nested snarls, within the chain being skipped, that the automaton's read head was is in before encountering the current symbol. Under that is a running distance along the chain being skipped, which no longer needs to be used. Under that are distances for each thing still to the left in the snarl containing the chain being skipped, including the snarl start. And under those are similar distances for higher levels of enclosing snarl that the automaton has read into.

When a SEED, EDGE, NODE_COUNT, or CHAIN_END is encountered, it is ignored.

When a SNARL_END is encountered, the nested snarl count at the top of the stack is increased by one, so that CHAIN_START within the child snarl can be recognized as nested and ignored.

When a SNARL_START is encountered, the nested snarl count at the top of the stack is decreased by one.

When a CHAIN_START is encountered, the automaton checks the count of nested snarls that it is currently reading through within the chain it is skipping, at the top of the stack. If it is not zero, the symbol is ignored, because the chain it belongs to must be nested somewhere inside a snarl along the chain that the automaton is skipping to the start of. If, on the other hand, it is zero, then the automaton knows it has reached the start of the chain it has been trying to skip, and can stop skipping. The automaton then pops both the number of nested snarls being traversed and the running distance (which was not updated after the decision to skip the chain was made) off of the stack, leaving the distances for other things in the skipped chain's parent snarl at the top, and goes to SCAN_SNARL.

##### S2.5.2 Processing enumerated source seeds

For each potential source seed for a given destination seed (see S2.5), we check if either the source and destination seeds are both on the forward strand of the read, or both on the reverse. If they are both on the forward strand, we find the anchor corresponding to the source seed (see S3), skipping the seed if there is no such anchor, and record a transition from the source anchor to the destination anchor with the zip code tree's estimated graph distance between the seeds. If they are both on the reverse strand, we find the anchor, but then *exchange the source and destination anchors* when recording the transition. This amounts to following the reverse strand through the graph along the path the zip code tree is measuring, to match the order and orientation of seeds in the read.

##### S2.5.3 Processing transitions

After iteration through the zip code tree from left to right is complete, all the recorded transitions are sorted in order of increasing anchor start position in the read. This handles multiple occurrences of entries for the same seed in the zip code tree, and allows for the read to wind back and forth through the snarls and chains modeled by the zip code tree without breaking the optimal substructure of the dynamic programming problem. Transitions are then filtered and used for chaining (see Section S3.1).

### S3 Chaining

The general problem of chaining starts with a set of **anchors** as input. An anchor is defined by an interval on the read and a corresponding interval on the reference.

A chain is an ordered subset of anchors that are co-linear in the read and reference. Chaining algorithms generally attempt to find an optimal chain that maximizes coverage over the read and minimizes the cost of gaps between consecutive anchors. An optimal solution to the chaining problem can be found in a simple dynamic programming recurrence over the anchors.

#### S3.1 Chaining in Giraffe

Giraffe performs two rounds of chaining to get the final chains of seeds. In the first round, we use seeds as anchors and produce chains of seeds which we call **fragments**. In the second round, we use the resulting fragments as anchors and perform the same chaining procedure, this time with a more permissive lookback distance, to produce the final **chains**. We found that two rounds of chaining reduced the time and memory use of chaining while still producing useful chains. Note that by having two rounds, we necessarily make greedy decisions at the first round, and we are not globally optimizing for the best chain over all the seeds.

To support having two rounds of chaining, Giraffe models anchors as starting and ending at seeds (which may be the same seed), and extending some distance beyond the seeds' outer endpoints.

We define the notion of the **corresponding anchor** for a seed, which depends on whether the seed is on the forward or reverse strand of the read. A seed may have no corresponding anchor in a given chaining problem, but we set up the problems so a seed never has more than one. If a seed is on the forward strand of the read, then as a source seed its corresponding anchor is the anchor that ends at the seed, and as a destination seed its corresponding anchor is the anchor that starts at the seed. If the seed is on the reverse strand, then as a source seed its corresponding anchor is the anchor that *starts* at the seed instead, and as a destination seed its corresponding anchor is the anchor that ends at the seed.

Anchors also have a **read exclusion zone** tracking the read intervals covered by their seeds' minimizers, to prevent chaining together seeds whose minimizers overlapped in the read (before they were trimmed down to single-graph-node seeds).

After the transitions between seeds from the zip code tree are sorted (see Section S2.5), they are filtered to remove transitions where the destination anchor is not actually reachable from the source anchor in the read, where the distance from the source anchor to the destination anchor in the read is longer than the **read lookback limit** (which is computed similarly to the graph lookback limit, but with different constants), or where the anchors impinge on one another's read exclusion zones (because their minimizers overlapped in the read).

The graph distance between the seeds is decreased by the lengths of the parts of their corresponding anchors that extend towards each other past the boundaries of their seeds. This gives the graph distance between the anchors. Then, the transition is discarded if this indicates that the anchors overlap in the graph.

Finally, the transition from the source anchor to the destination anchor, with its associated read and graph distances, is used to drive the dynamic programming algorithm for chaining.

To score chains, Giraffe uses a scoring scheme based on that of Minimap2 [10]. The overall score of a chain is the sum of the scores of the anchors and the scores of the gaps between consecutive anchors. The score of an anchor is its alignment score, scaled by a user-defined factor, plus a user-defined bonus score (Section S3.2). The cost of a gap between two consecutive anchors is found from the absolute difference (*dist*) between the graph distance (from the zip code tree) and the read distance, if it is nonzero, and the length (*seed_length*) of the minimizers used for seeding. This is the same gap cost function as is used in Minimap2, near equation 2 of Li [10]:

$$gap\_cost = 0.01 * seed\_length * dist + 0.5 * log_2(dist) \quad (1)$$

Giraffe also has the option to consider the matches that would be taken in the gap between two consecutive anchors. The number of matches, which is the minimum of the distances between the anchors in the read and the graph, is scaled by a user-defined factor specifying the number of points per potential match in the gap (Section S3.2). The overall score of a gap is therefore:

$$gap\_score = matches * scale\_factor - gap\_cost \quad (2)$$

#### S3.2 Fragment and chaining parameters

When doing fragmenting, we use a lookback distance that is the maximum of the *fragment-max-lookback-bases* and the read length times the *fragment-max-lookback-bases-per-base*. We also limit the indel length for fragmenting to be the maximum of the *fragment-max-indel-bases* and the read length times the *fragment-max-indel-bases-per-base*. Giraffe will make at most *max-fragments* fragments total. The fragments are scored based on the gap costs between consecutive pairs of seeds. We scale the gap score by a factor of *fragment-gap-scale*. We add *fragment-points-per-possible-match* to the score for non-indel connecting bases. We add *item-bonus* to the score for each seed included in the fragment and the score of each seed is scaled by a factor of *item-scale*.

We discard fragments if their score is not good enough relative to the score of the best-scoring fragment. We set the score limit to be the best score times the *fragment-score-fraction*, as long as this value is between the *fragment-min-score* and the *fragment-max-min-score*. If this value is greater than *fragment-max-min-score* or less than *fragment-min-score*, it is set to the bounding value. A fragment is also discarded if the zip code tree that it came from was less than *fragment-set-score-threshold* away from the score of the best tree. We will always keep at least *min-chaining-problems* and at most *max-chaining-problems* fragments.

We use similar parameters for doing chaining over the fragments. The lookback distance for chaining is the maximum of the *max-lookback-bases* and the read length times the *max-lookback-bases-per-base*. The maximum indel length is the maximum of the *max-indel-bases* and the read length times the *max-indel-bases-per-base*. The gap score is scaled by a factor of *gap-scale*. We add *item-bonus* to the score for each fragment included in the chain, and the score of each fragment is scaled by a factor of *item-scale*. We add *points-per-possible-match* for each non-indel connecting base in the chain.

A chain is only kept if its score is within *chain-score-threshold* of the score of the best-scoring chain. The score of a chain must also be higher than the minimum of the *max-min-chain-score* and the length of the read times the *min-chain-score-per-base*. Regardless of score, at least *min-chains* are kept. We keep up to *max-chains-per-tree* chains that originated from the same zip code tree.

### S4 Aligning

To extend a chain of seeds into an alignment, Giraffe performs two types of base-level alignment: alignment between consecutive pairs of seeds in a chain and pinned tail alignment anchored on the first or last seed in a chain. For both types of alignment, we first try to use a wavefront alignment (WFA) algorithm [53, 55] to align to the haplotypes in the GBWT. If the WFA extension approach is unsuccessful for a between-seed alignment (because it hits a score penalty limit, for example, or because the problem is too large), then we use a banded global aligner to align against the graph itself. If the problem is also too big for the banded global aligner, then we treat the first seed in the pair as the end of the chain and do pinned tail alignment from this seed. For tail alignments that can't be done with the WFA aligner, we use Dozeu, a SIMD-accelerated X-drop aligner.

The final alignment is strongly influenced by the chain used to find it. For example, if the chain contains a seed in an insertion node, then the final alignment is forced to take that insertion, even if a more optimal alignment could be found without the seed. This case is common in repetitive regions where seeds with identical sequences are found near each other in the reference graph. To avoid forcing the alignment to include a repetitive seed, we flag seeds as being in repetitive regions of the read and allow the aligner to skip repetitive seeds in a chain.

Seeds are flagged as being in a repetitive region using a Hidden Markov Model (HMM). The seeds are first ordered according to their position in the read. Each seed will be labeled as having a hidden state of "unique" or "repetitive" based on the number of occurrences of the seed's minimizer in the graph. The probability of switching between the states is 0.1 and the probability of emitting an occurrence count of 1 with a hidden state of "repetitive" (or an occurrence count greater than 1 with a hidden state of "unique") is 0.1. The

HMM is initialized with a 0.05 probability of being repetitive.

The repetitive flag is not taken into account for chaining the seeds or for scoring the chains. When aligning between seeds in the chain, each consecutive repetitive seed in sequence along a chain will be added to a candidate skip set, as long as the sum of the graph distances into, between, and out of the sequence of seeds is below *max-skipped-bases*. Then the whole set will be skipped if the sum of absolute gap sizes (the difference between the distance in the read and the distance in the graph) required in the seed connections being skipped is more than 50 bp. The first and last seeds in a chain are never skipped.

#### S4.1 Alignment parameters

To avoid using too many resources during alignment, we set limits on the sizes of problems that each aligner can attempt. When aligning either tails and between-seed connections with the WFA aligner, we limit the number of mismatches or equivalent-scoring gaps based on the length of the read. The number of mismatches is capped at the smaller of *wfa-max-max-mismatches* and the *wfa-max-mismatches* plus the read length times the *wfa-max-mismatches-per-base*. Similarly, band distance of WFA alignment is capped at the smaller of *wfa-max-distance* and the *wfa-distance* plus the read length times the *wfa-distance-per-base*.

When aligning between seeds in a chain, we will only connect seeds across a maximum distance of *max-chain-connection* with WFA. If the length of the gap between seeds in the read is greater than *max-chain-connection* but less than *max-middle-dp-length*, then we align it with the banded global aligner. If it is longer than *max-middle-dp-length*, then we make it a tail alignment and attempt to align it using Dozeu. When skipping repetitive seeds in a chain, seeds may only be skipped if the gap introduced will be less than *max-skipped-bases*.

When aligning tails, we only attempt alignment if the aligner will use fewer than *max-dp-cells* cells in the dynamic programming matrix. If the length of the tail is less than *max-tail-length*, then we will attempt to use the WFA extender. If the tail is longer than the *max-tail-length* but shorter than *max-tail-dp-length*, then we do the alignment with Dozeu. For tails that are longer than *max-tail-dp-length*, we make a softclip. When aligning a tail, we allow at most *max-tail-gap* gap bases.

#### S4.2 Surjection

For variant calling with a linear-reference caller like DeepVariant, aligned long reads needed to be re-expressed in the space of, and partially re-aligned to, the linear reference. This process is called **surjection**, and is a longstanding feature of the `vg` toolkit. However, to achieve good variant calling from long reads, the surjection algorithm in `vg` had to be modified.

The surjection algorithm works by finding **anchors** where the read already touches nodes that lie along the target linear reference's path through the graph. (These are similar in concept to, but distinct from, the seed-based anchors used in chaining during long read alignment.) The algorithm already had a facility for **pruning** anchors in regions of low sequence complexity (which have a tendency to produce high-complexity graphs). In these regions, errors in the alignment of the read to the graph, and of the alignment within the graph between the different haplotypes it represents, are more likely, because it's hard to tell where you are by looking at the sequence. Alignment errors can compound when a match against a node in the graph is treated as a match against a linear path that visits that node: if either input match assertion was wrong, the output will be wrong. To deal with this, the anchor pruning logic removes anchors consisting of low-complexity sequence. It is activated by the `--prune-low-cplx` option to `vg surject`, which we used in our surjection operations. Finally, after pruning the anchors, `vg surject` aligns the portions between them, and the tails, to the linear reference. This is somewhat like how Giraffe extends a chain into a final graph alignment.

Some improvements in long read surjection came from changes to Dozeu and other components shared with Giraffe. We also added giving-up logic to detect and skip intractably large alignment problems, to make sure long read surjection was fast enough to be practical. However, we found that we also had to supplement the existing anchor pruning logic to get good DeepVariant calling results.

The existing filter was based on a statistical analysis of anchor sequence complexity. We added a **simple slide** pruning algorithm on top of this. If a proposed anchor between the read and the linear reference and the read re-occurs in the read sequence within (by default) 6 bp (or twice the anchor length, whichever is lower), we prune the anchor away. This is designed to allow surjection to second-guess the graph's alignment decisions for homopolymers and short repeats, and to produce alignments of these regions against the linear reference that are more consistent between different reads with slightly different sequences. (We had noticed that a single base added or dropped in a homopolymer, or a single base error in a repeat unit, could cause a read to take a very different optimal local path through the graph, resulting in a very different surjection that would then confuse downstream tools.) If, in the future, a long-read-capable linear-space indel realignment tool, or a graph-space indel realignment tool, were developed, this additional pruning step might prove unnecessary.

### S5 Evaluation methods

#### S5.1 Graphs

For the linear mappers, we mapped to the CHM13 reference genome [59]. We also used Giraffe to map to a linear graph containing only CHM13.

The graph references used were constructed from the HPRC release 2 assemblies using Minigraph and Minigraph-Cactus using CHM13 as the reference. We used graphs that were not built with the HG002 sample we used for evaluation, which are distinguished in their filenames as "eval" graphs; real applications should use non-"eval" graphs that include HG002 and the Y chromosome it contributes to the CHM13v2.0 linear reference. The HPRC frequency filtered graph (HPRC-d46) was constructed using the Minigraph-Cactus pipeline, with an allele frequency filtering threshold of 46. These graphs, along with instructions to reproduce them, are currently available here: `https://s3-us-west-2.amazonaws.com/human-pangenomics/index.html?prefix=pangenomes/scratch/2025_02_28_minigraph_cactus/`. Note that the `hprc-v2.prerelease.3-mc-chm13-eval-jan29` graph used for training was produced with an earlier version of the pipeline, `https://github.com/ComparativeGenomicsToolkit/cactus/commit/c1c366ceb519c85bc2bc50e8f1d703b1`[illegible] using an earlier version of the assemblies, `https://github.com/human-pangenomics/hprc_intermediate_assembly/blob/b39df38971b4cf138421b2f1896dde798c221803/data_tables/assemblies_pre_release_v0.4.index.csv` as input. This graph is not available as it has been deprecated in favor of the newer graphs. The HPRC sample graphs (HPRC-Sampled16) were constructed with the Minigraph-Cactus selectively clipped graph (HPRC-clipped) as the base. Each sampled graph was built using reads from the sample-and-sequencing-technology combination it was used with. (We only evaluate performance on HG002, but sampled graphs for other samples were produced during DeepVariant model training in S5.4.1.) Sampling was done using real read sets, and the same sampled graph was used for mapping both real and simulated reads.

In addition to the graphs constructed with the HPRC release 2 data, we also mapped to graphs from the HPRC release 1. The HPRC-v1-Sampled32 graph was constructed using the haplotype sampling pipeline with 32 haplotypes from the clipped, non-frequency filtered, CHM13-based HPRC v1.1 graph available from `https://github.com/human-pangenomics/hpp_pangenome_resources`.

#### S5.2 Reads

We used real reads from HG002 [60] sequenced with PacBio's HiFi, Oxford Nanopre's R10 chemistry, and Illumina's NovaSeq, and with an Element Biosciences instrument. We used these reads to perform variant calling and to assess the speed, memory use, and number of softclipped, hardclipped, and unmapped bases of each mapping tool.

HiFi reads for HG002 were provided by Aaron Wenger at PacBio to the Telomere-to-Telomere Consortium for the "Q100" benchmark project described at `https://github.com/marbl/HG002` (and more specifically `https://github.com/marbl/HG002/blob/main/Sequencing_data.md#hifi-data`), and were retrieved from `https://s3-us-west-2.amazonaws.com/human-pangenomics/index.html?prefix=T2T/HG002/assemblies/polishing/HG002/v1.0/mapping/hifi_revio_pbmay24/`.

R10 reads for HG002 were selected specifically to be from recent sequencing runs in 2025 and later, because we noticed that current R10 read characteristics differed noticeably from runs from earlier in the history of the chemistry. Internally in the analysis scripts, we refer to this as `r10y2025` data, as distinct from older `r10`, though we tuned the Giraffe `r10` preset for this regime. The data used came from Oxford Nanopore's "Genome in a Bottle Data Release 2025.01", documented at https://epi2me.nanoporetech.com/giab-2025.01/, and downloaded from s3://ont-open-data/giab_2025.01/.

Illumina NovaSeq reads for HG002 were retrieved from Google's gold-standard benchmarking dataset collection [61] at gs://brain-genomics-public/research/sequencing/fastq/novaseq/wgs_pcr_free/40x/.

Element reads for HG002 were retrieved from https://s3-us-west-2.amazonaws.com/human-pangenomics/index.html?prefix=T2T/scratch/HG002/sequencing/element/trio/HG002/ins500_600/.

Sets of one million reads were simulated with PBSIM2 [62] from the HG002 sample, which was not included in any of the reference graphs, for each sequencing technology (see Section S5.2). Long reads were simulated using a graph built with Minigraph-Cactus using the HG002 assembly, GRCh38, and CHM13 as input. The HG002 assembly was extracted from the graph, then PBSIM2 was used to simulate reads. PBSIM2 was run with parameters `--depth 4 --accuracy-min 0.00 --length-min 10000 --difference-ratio 6:50:54` for long reads, and `--depth 1 --accuracy-min 0.00 --length-min 250 --length-max 250 --difference-ratio 7:1:1` for Illumina reads. The command also used the `--hmm_model` feature to simulate reads based on real quality strings, using our real HG002 read sets for each sequencing technology. The simulated reads were then converted to BAM format and projected onto the HG002 graph with `vg inject`. The reads were then annotated with the reference positions that they overlapped in the graph with `vg annotate`. Element reads were simulated using `vg sim` with parameters `--frag-len 550 --frag-std-dev 100 --sub-rate 0.0003 --indel-rate 0.00000001`, based on error rates from Liu et al. [63], using real reads from the HPRC consortium available at https://s3-us-west-2.amazonaws.com/human-pangenomics/index.html?prefix=T2T/scratch/HG002/sequencing/element/trio/HG002/ins500_600/.

#### S5.3 Mapping experiments

We compared Giraffe to the linear mappers Minimap2 version 2.28-r1209 [10], Winnowmap version 2.03 [34], and BWA-MEM version 0.7.18-r1243 [9], as well as the graph mappers GraphAligner version 1.0.20 [32], and Minigraph version 0.21-r606 [26]. GraphAligner was run using a container available at quay.io/biocontainers/graphaligner:1.0.20--h06902ac_0 and Giraffe was run on commit `99930d` for all experiments.

Unless otherwise specified, we ran Minimap2 with presets `lr:hqae` for R10 reads, `map-hifi` for HiFi reads, and `sr` for Illumina reads. We also tested the presets `map-ont` for R10 reads and `map-pb` and `lr:hqae` for HiFi reads, but we found that `map-hifi` was both faster and more accurate than the other presets (Table S3), and `lr:hqae` was more than twice as fast and only slightly less accurate (Tables S4). The runtimes of these parameters were found using different read sets than other results included in this paper and are therefore not shown.

Giraffe was run with parameter presets `hifi` and `r10` for HiFi and R10 reads respectively. For HiFi reads, Giraffe was provided the `--max-min-chain-score 100` option (see Section S3.2). As of vg version 1.68.0, this parameter change is incorporated into the base `hifi` preset. Giraffe used minimizers constructed using `vg minimizer`, using different parameters for short and long reads. For short reads, minimizers were constructed using options `--kmer-length 29 --window-length 11`; for long reads, minimizers were constructed using options `--kmer-length 31 --window-length 50 --weighted`. In current versions of `vg`, these are now the default settings for the `sr-giraffe` and `lr-giraffe` workflow targets for `vg autoindex`, respectively.

The `vg` version used for surjection was not controlled in our evaluation pipeline for mapping and calling.

Mapping correctness on simulated reads was determined using the reference annotations that were added when simulating the reads (Section S5.2). For linear mappers, the reads were mapped onto the linear reference genome and then projected onto the HPRC-filtered graph using `vg inject`. Read-to-graph alignments were annotated with reference positions using `vg annotate -m --search-limit=-1`. Correctness was found using `vg gamcompare --range 200`. This means that if an annotation on the read was within 200 bp of any reference annotation on the simulated read, then the read was counted as being correctly mapped.

We note that some reads could not be annotated with reference coordinates. This may occur in regions of the genome where the sample differes greatly with the reference. Reads with no reference coordinates are labeled as "ineligible", as the correctness of a mapping cannot be determined. These reads are therefore are not included in the ROC plots, however they are included in the overall mapping accuracy tables.

GraphAligner was run using the default parameters and also using a parameter set recommended by the developer to improve the speed. The `GraphAligner (fast)` setting used the following parameters: `--seeds-mxm-length 30 --seeds-mem-count 10000 --bandwidth 15 --multimap-score-fraction 0.99 --precise-clipping 0.85 --min-alignment-score 100 --clip-ambiguous-ends 100 --overlap-incompatible-cutoff 0.15 --max-trace-count 5 --mem-index-no-wavelet-tree`.

All analyses were run using Snakemake on a Slurm cluster, using a Ceph-based shared filesystem for data storage. Analyses being timed were restricted to identical nodes with 2 TB of memory, 14 TB of NVMe scratch (plus the system drive), and two AMD EPYC 7662 CPUs, providing:

- 2 sockets
- 128 total physical cores
- 256 total logical cores, or hardware threads
- 32 KiB L1 instruction cache and 32 KiB L1 data cache per physical core
- 512 KiB L2 cache per physical core
- 512 MiB total L3 cache

All cores were the same type, implemented using the same microarchitecture. Simultaneous Multithreading (SMT) was left enabled, allowing each physical core to provide two logical cores. Frequency boost was left enabled, allowing CPU cores to boost above their base clock speed of 2 GHz.

For accurately finding the runtime and memory use of each mapper, we ran mapping jobs being timed jobs with the `--exclusive` flag to ensure that they were run on a node with no other running jobs. All mappers were run with 64 threads. We were unable to finish running GraphAligner with default parameters on the HPRC-d46 graph with the whole HiFi or R10 read sets, so real read statistics including runtime and memory use that are comparable to the other mappers are not available.

#### S5.4 Short variant calling with DeepVariant

##### S5.4.1 DeepVariant training

For HiFi reads, we trained a DeepVariant model, called the `2025-03-26noinfo` model, on Giraffe alignments of HiFi reads.

Read files used were obtained from:

- The five non-"analysis" BAM files from `https://downloads.pacbcloud.com/public/revio/2022Q4/`.
- `https://storage.googleapis.com/brain-genomics/awcarroll/pacbio_training_2024/fastq/revio/NA12878.GRCh38.haplotagged.35x.fastq.gz`
- `https://storage.googleapis.com/brain-genomics/awcarroll/pacbio_training_2024/fastq/revio/NA12878.GRCh38.haplotagged.fastq.gz`
- Two amplified Revio "Twist cells" data files, for HG001 (at about 32x coverage) and HG002 (at about 33x coverage), and one amplified "ULI" data file for HG002 (at about 60x coverage), obtained from PacBio. These files are available from Google at `gs://brain-genomics-public/publications/chang2025_long_read_giraffe/`.
- Three PacBio Sequel2 files, for HG002, HG003, and HG004, from the input data for the precisionFDA Truth Challenge v2 [64]. These reads are available in SRA under BioProjects `PRJNA586863`, `PRJNA626365`, and `PRJNA626366`,
- Six PacBio SPRQ files, with run identifiers `m84029_240730_192004_s1` and `m84034_240730_194815_s1` (for HG002), `m84028_240802_214905_s1` and `m84029_240730_212142_s2` (for HG003), and `m84030_240730_192221_s1` and `m84030_240802_210346_s1` (for HG004), obtained from PacBio. These files are available from Google at `gs://brain-genomics-public/publications/chang2025_long_read_giraffe/`.

The training data was created by aligning these reads using vg commit ceb8ad76e537d66433f3078848489026709e7557, against haplotype-sampled versions of the hprc-v2.prerelease.3-mc-chm13-eval-jan29 graph, and then surjecting the reads to the GRCh38 linear reference. Alignment and surjection was orchestrated using commit 32af08d460f0985e373bc61f1c9f6ee52134f536 of the Giraffe workflow from the vg_wdl project (https://github.com/vgteam/vg_wdl), and the hifi-training.sbatch script at commit c547fff7c5385566a2493ded4d7afc84c803606a in the long-read-giraffe-experiments repository (https://github.com/vgteam/long-read-giraffe-experiments).

DeepVariant was trained using these reads and a truth set for each sample, using the Google-internal DeepVariant training process. No model.example_info.json file was added to the model, which would be required for usage with newer versions of DeepVariant.

Both the trained model and a version of the model which *does* have a model.example_info.json file, suitable for use with current releases of DeepVariant, are available on Zenodo [56].

##### S5.4.2 DeepVariant calling

Variant calling with DeepVariant was performed with commit 890d76647259aa412102dac412dd56f808e92f23, now tag lr-giraffe-paper, of the DeepVariant WDL workflow from the vg_wdl project, now available on Dockstore [65]. The workflow was run with several versions of Toil [66] sharing a WDL call cache (most recently commit 014a57d33cd34496f44006d09d4189f2d0ea3b9b based on the Toil 9.0.0 release), through the call_variants_dv rule in our Snakefile. This workflow was used to run a specially-prepared head756846963 version of DeepVariant, which is now published on Docker Hub as google/deepvariant:lr-giraffe-paper and google/deepvariant:lr-giraffe-paper-gpu. This version has the features of the DeepVariant 1.9 release, along with, most prominently, a bugfix to enable DeepVariant's --normalize_reads feature to work on long reads. For calling on HiFi data, a specially-trained DeepVariant model was used (see Section S5.4.1). For other sequencing technologies, the default models included with DeepVariant were used.

In the WDL workflow input parameters, read left-alignment and DeepVariant read normalization were turned on, but indel realignment was turned off (as we could not locate an indel realignment tool supporting long reads to add to the workflow). For R10 reads, the --small_model_call_multiallelics=false parameter was provided to DeepVariant via the DeepVariant.OTHER_MAKEEXAMPLES_ARG workflow parameter field, which turned off the "small model" fast-path calling approach for multiallelic variants. The workflow was provided with the list of expected-haploid contigs for the sample (chrX and chrY), as well as a BED file giving the locations of the pseudoautosomal regions in the target linear reference assembly. Single-BAM and calling-BAM output were turned off. 24 cores and 200 gigabytes of memory were allocated to calling tasks, and 50 gigabytes of memory were allocated to DeepVariant make_examples tasks.

This workflow was also used to evaluate calling accuracy; the standalone vcfeval run was turned off, because the VCFs used were sometimes able to cause vcfeval to crash. In addition to the high-confidence-region BED file for the variant truth set being used, the workflow was provided with a "restrict regions" BED file describing the regions in the target linear reference assembly that were actually present in the HPRC graphs being tested. (This served to exclude the Y chromosome of CHM13 from the calling accuracy analysis. CHM13 itself does not carry a Y chromosome. The CHM13v2.0 linear reference, where the benchmark VCF is described, uses a Y chromosome sequence that actually came from the HG002 donor. But the "eval" HPRC graphs held out the HG002 sample so it could be used as a test set, and used a different Y sequence in their "CHM13" reference. To work around this, CHM13's Y was excluded from the DeepVariant calling analysis.)

##### S5.5 Structural variant calling and genotyping

Structural variant genotyping was run using a pipeline similar to that of Hickey et al. [41]. The coverage of the read alignments on the graph was computed using `vg pack` (options `--min-mapq 5 --with-edits`). Variants were called from the resulting read coverage file (options `--all-snarls --gbz --min-length 30`). We then used `vcfbub` with options `-l 0 -a 100000` and `vcfwave` with options `-I 1000` to decompose nested variants [67].

For structural variant calling against a linear genome, we used Sniffles version 2.5.3 [42, 43]. Sniffles was run with tandem repeat annotations from Pacific Biosciences `https://github.com/PacificBiosciences/pbsv/blob/master/annotations/human_chm13v2.0_maskedY_rCRS.trf.bed`.

We ran these structural variant calling and genotyping pipelines on the same HG002 reads that were used for small variant calling. We compared our structural variant calling and genotyping results to the Genome in a Bottle Consortium's `CHM13v2.0_HG2-T2TQ100-V1.1` truth set [44]. The truth VCF and BED files were accessed from `https://ftp-trace.ncbi.nlm.nih.gov/ReferenceSamples/giab/data/AshkenazimTrio/analysis/NIST_HG002_DraftBenchmark_defrabbV0.018-20240716/`.

We then used Truvari [45] version 4.3.1 to benchmark the structural variant calls from `vg call` and Sniffles. Benchmarking was done using `truvari bench` with options `--pick ac --passonly -r 2000 -C 5000`, and `truvari refine` with options `--recount --use-region-coords --use-original-vcfs --align mafft/can`. Benchmarking was restricted to regions specified by the `svvar` BED file provided by GIAB.

#### S6 Pangenome-guided assembly

##### S6.1 Reads

For pangenome-guided assembly testing, we used HG002 nanopore R10 reads (sample PAW70337), released by GIAB at `https://epi2me.nanoporetech.`

`com/giab-2025.01/` (and more specifically `s3://ont-open-data/giab_2025.01/basecalling/sup/HG002/PAW70337/`). Pangenome-guided assembler (PGA) demonstration for the assembly of RCCX haplotypes used two Congenital Adrenal Hyperplasia (CAH) samples from the Disorders of Sex/Development—Translational Research Network (DSD-TRN) biobank, which were sequenced at UCSC and analyzed as part of this work [46, 47]. These data are not publicly released but are available from the authors upon request.

##### S6.2 RCCX locus

The RCCX locus is a complex, multiallelic tandem copy-number variation on chromosome 6p21.3, located within the major histocompatibility complex class III region. It comprises one or more repeating modules, each containing four closely linked genes: STK19 (serine/threonine kinase 19), C4 (complement component 4), CYP21 (steroid 21-hydroxylase), and TNX (tenascin-X) ([68, 69]).

Pathogenic variants and rearrangements of *CYP21A2* are the primary cause of Congenital Adrenal Hyperplasia (CAH), an autosomal recessive disorder (OMIM #201910). The module also contains *CYP21A1P*, a highly homologous pseudogene sharing 99.6% coding-sequence identity with *CYP21A2*. The high sequence similarity at this locus predisposes it to frequent non-allelic homologous recombination, which, in rare instances, can be deleterious, accounting for over 95% of CAH cases. These recombination events occur either as large deletions or as gene conversions. Deletions, typically caused by unequal crossover, produce non-functional chimeric *CYP21A1P/CYP21A2* genes and account for approximately 30% of cases. Gene conversions, in which deleterious variants from the pseudogene are passed to the active gene, explain about 65% of cases [70].

Beyond CAH, the RCCX locus is clinically significant because it also harbors the *TNXB* gene (where biallelic variants are associated with Ehlers-Danlos syndrome, classic-like, 1, OMIM #606408)([71, 72]), and the *C4* gene (where copy number variations are associated with susceptibility to schizophrenia and autoimmune conditions) [73]. The region's structural complexity is further compounded by a truncated *TNXA* pseudogene and a polymorphic HERV-K insertion, making it particularly challenging to resolve with standard genomic methods ([74]).

#### S7 Supplemental Tables

| Reference | Mapper | Correct (%) | Mapq60 (%) | Wrong Mapq60 (%) |
|---|---|---|---|---|
| CHM13 | BWA-MEM | 92.1348 | 89.9381 | 0.0493 |
| CHM13 | Giraffe | 92.0959 | 90.3686 | 0.0551 |
| CHM13 | Minimap2-sr | 91.9596 | 89.3215 | 0.0678 |
| HPRC-d46 | Giraffe | 92.0825 | 90.0379 | 0.0343 |
| HPRC-sampled16 | Giraffe | 91.9942 | 89.9318 | 0.0312 |
| HPRC-v1-Sampled32d | Giraffe | 92.136 | 90.2477 | 0.0398 |

Table S1: **Illumina mapping accuracy**. Mapping correctness and mapping quality statistics for 1 million simulated Illumina reads.

| Reference | Mapper | Correct (%) | Mapq60 (%) | Wrong Mapq60 (%) |
|---|---|---|---|---|
| CHM13 | BWA-MEM | 92.1253 | 89.8975 | 0.0498 |
| CHM13 | Giraffe | 92.1129 | 90.6621 | 0.0621 |
| CHM13 | Minimap2-sr | 92.0627 | 89.7795 | 0.0557 |
| HPRC-d46 | Giraffe | 92.0702 | 90.2396 | 0.0367 |
| HPRC-sampled16 | Giraffe | 91.9993 | 90.0879 | 0.0333 |
| HPRC-v1-Sampled32d | Giraffe | 92.1431 | 90.4701 | 0.0476 |

Table S2: **Element mapping accuracy**. Mapping correctness and mapping quality statistics for 1 million simulated Element reads.

| Reference | Mapper | Correct (%) | Mapq60 (%) | Wrong Mapq60 (%) |
|---|---|---|---|---|
| CHM13 | Giraffe | 92.4807 | 89.7735 | 0.0299 |
| CHM13 | Minimap2-lr:hqae | 92.4911 | 93.6124 | 0.215 |
| CHM13 | Minimap2-map-hifi | 92.5093 | 92.9463 | 0.1697 |
| CHM13 | Minimap2-map-pb | 92.5066 | 93.1879 | 0.1731 |
| CHM13 | PBMM2 | 92.5086 | 93.1188 | 0.1749 |
| CHM13 | Winnowmap | 92.5006 | 97.0862 | 0.5031 |
| HPRC-d46 | Giraffe | 92.6419 | 90.4139 | 0.0199 |
| HPRC-d46 | GraphAligner-default | 92.582 | 99.1285 | 0.5209 |
| HPRC-d46 | GraphAligner-fast | 92.5636 | 98.9838 | 0.5308 |
| HPRC-Minigraph | GraphAligner-default | 92.0684 | 99.2341 | 1.0319 |
| HPRC-Minigraph | GraphAligner-fast | 92.113 | 99.0453 | 0.9781 |
| HPRC-Minigraph | Minigraph | 91.1538 | 89.9867 | 0.647 |
| HPRC-Sampled16 | Giraffe | 92.5853 | 90.3469 | 0.0171 |
| HPRC-v1-Sampled32d | Giraffe | 92.594 | 90.5621 | 0.0227 |

Table S3: **HiFi mapping accuracy**. Mapping correctness and mapping quality statistics for 1 million simulated PacBio HiFi reads.

| Reference | Mapper | Correct (%) | Mapq60 (%) | Wrong Mapq60 (%) |
|---|---|---|---|---|
| CHM13 | Giraffe | 91.8138 | 93.1543 | 0.27 |
| CHM13 | Minimap2-lr:hqae | 91.981 | 92.2521 | 0.2053 |
| CHM13 | Minimap2-map-ont | 92.4468 | 92.2967 | 0.1541 |
| CHM13 | Winnowmap | 92.2549 | 94.3931 | 0.3374 |
| HPRC-d46 | Giraffe | 92.02 | 92.8973 | 0.1468 |
| HPRC-d46 | GraphAligner-default | 92.4183 | 98.5867 | 0.5767 |
| HPRC-d46 | GraphAligner-fast | 92.2038 | 98.123 | 0.6212 |
| HPRC-Minigraph | GraphAligner-default | 91.7993 | 98.615 | 1.1444 |
| HPRC-Minigraph | GraphAligner-fast | 90.694 | 98.0489 | 2.0635 |
| HPRC-Minigraph | Minigraph | 90.5213 | 89.3673 | 1.0062 |
| HPRC-Sampled16 | Giraffe | 91.971 | 92.7292 | 0.1536 |
| HPRC-v1-Sampled32d | Giraffe | 91.9936 | 92.9659 | 0.1729 |

Table S4: **R10 mapping accuracy**. Mapping correctness and mapping quality statistics for 1 million simulated Oxford Nanopore R10 reads.

| Reference | Mapper | Runtime (min) | Index time (min) (+ Sampling (min)) | Memory (GB) | Unmapped Bases (%) |
|---|---|---|---|---|---|
| CHM13 | BWA-MEM | 245.6833 | 0.0000 | 18.5305 | 0.1831 |
| CHM13 | Giraffe | 79.5333 | 1.2549 | 45.3265 | 0.9384 |
| CHM13 | Minimap2-sr | 117.1167 | 0.2639 | 13.7634 | 0.3192 |
| HPRC-d46 | Giraffe | 144.0833 | 2.2578 | 68.3341 | 0.9080 |
| HPRC-sampled16 | Giraffe | 136.6500 | 1.6847 (+8.883) | 58.1731 | 0.9130 |
| HPRC-v1-Sampled32d | Giraffe | 110.0833 | 1.5023 (+8.65) | 52.3584 | 0.8240 |

Table S5: **Mapping statistics for real Illumina reads**. Real Illumina reads were mapped to various linear and graph references. Giraffe was run against the graphs directly, or using a haplotype sampling pipeline sampling either 16 haplotypes, or 2 haplotypes from 32 in diploid mode. The time for haplotype sampling is reported separately from the time for index loading and building. BWA-MEM does not report the time used for indexing. Runtime includes the time for indexing and haplotype sampling.

| Reference | Mapper | Runtime (min) | Index time (min) (+ Sampling (min)) | Memory (GB) | Unmapped Bases (%) |
|---|---|---|---|---|---|
| CHM13 | BWA-MEM | 335.2500 | 0.0000 | 21.4655 | 0.0742 |
| CHM13 | Giraffe | 112.8333 | 1.1626 | 45.3730 | 0.8468 |
| CHM13 | Minimap2-sr | 145.5333 | 0.2946 | 14.1044 | 0.1423 |
| HPRC-d46 | Giraffe | 190.4500 | 2.2604 | 68.5315 | 0.8279 |
| HPRC-sampled16 | Giraffe | 196.3500 | 1.7078 (+10.2333) | 58.6470 | 0.8322 |
| HPRC-v1-Sampled32d | Giraffe | 161.3000 | 1.4330 (+7.7833) | 52.5898 | 0.7435 |

Table S6: **Mapping statistics for real Element reads**. Real Element reads were mapped to various linear and graph references. Giraffe was run against the graphs directly, or using a haplotype sampling pipeline sampling either 16 haplotypes, or 2 haplotypes from 32 in diploid mode. The time for haplotype sampling is reported separately from the time for index loading and building. BWA-MEM does not report the time used for indexing. Runtime includes the time for indexing and haplotype sampling.

| Reference | Mapper | Runtime (min) | Index time (min) (+ Sampling (min)) | Memory (GB) | Soft- or Hard-clipped Bases (%) | Unmapped Bases (%) |
|---|---|---|---|---|---|---|
| CHM13 | Giraffe | 99.6333 | 0.3958 | 40.1394 | 0.1464 | 0.0307 |
| CHM13 | Minimap2-map-hifi | 84.1667 | 0.2347 | 35.6159 | 0.2311 | 0.0233 |
| CHM13 | PBMM2 | 164.4167 | NA | 44.9850 | 0.2147 | 0.0211 |
| CHM13 | Winnowmap | 980.7833 | 2.7004 | 84.5537 | 0.2282 | 0.0215 |
| HPRC-d46 | Giraffe | 115.2833 | 1.5775 | 68.9414 | 0.1604 | 0.0271 |
| HPRC-d46 | GraphAligner-fast | 2779.7500 | NA | 100.3282 | 0.8447 | 0.0210 |
| HPRC-Minigraph | Minigraph | 308.7833 | 2.6168 | 47.3127 | 2.1550 | 2.3308 |
| HPRC-sampled16 | Giraffe | 127.6667 | 0.9452 (+ 22.0333) | 69.3227 | 0.1384 | 0.0255 |
| HPRC-v1-Sampled32d | Giraffe | 110.4167 | 0.7522 (+ 24.65) | 53.9039 | 0.1060 | 0.0256 |

Table S7: **Mapping statistics for real HiFi reads**. Real HiFi reads were mapped to various linear and graph references. Minimap2 was run with the `map-hifi` setting. Giraffe was run against the graphs directly, or using a haplotype sampling pipeline sampling either 16 haplotypes, or 2 haplotypes from 32 in diploid mode. The time for haplotype sampling is reported separately from the time for index loading and building. GraphAligner does not report the time used for indexing. Runtime includes the time for indexing and haplotype sampling.

| Reference | Mapper | Runtime (min) | Index time (min) (+ Sampling (min)) | Memory (GB) | Soft- or Hard-clipped Bases (%) | Unmapped Bases (%) |
|---|---|---|---|---|---|---|
| CHM13 | Giraffe | 172.9667 | 0.3848 | 101.0554 | 2.7481 | 2.6964 |
| CHM13 | Minimap2-lr:hqae | 168.7500 | 0.0889 | 29.4945 | 4.1112 | 0.3345 |
| CHM13 | Winnowmap | 813.9167 | 2.7289 | 83.6441 | 3.5437 | 0.1648 |
| HPRC-d46 | Giraffe | 151.7500 | 1.5268 | 166.5571 | 2.7155 | 2.7986 |
| HPRC-d46 | GraphAligner-fast | 1557.4833 | NA | 100.3363 | 7.2328 | 0.1501 |
| HPRC-Minigraph | GraphAligner-default | 8404.4500 | NA | 49.5475 | 4.3271 | 0.0057 |
| HPRC-Minigraph | Minigraph | 404.8167 | 2.6236 | 53.9274 | 5.6931 | 2.5676 |
| HPRC-sampled16 | Giraffe | 143.3833 | 0.9395 (+ 23.233) | 137.2166 | 2.7065 | 2.7325 |
| HPRC-v1-Sampled32d | Giraffe | 127.1500 | 0.7654 (+ 22.1) | 142.5726 | 2.7110 | 2.8703 |

Table S8: **Mapping statistics for real R10 reads**. Real R10 reads were mapped to various linear and graph references. Minimap2 was run with the `lr:hqae`setting. Giraffe was run against the graphs directly, or using a haplotype sampling pipeline sampling either 16 haplotypes, or 2 haplotypes from 32 in diploid mode. The time for haplotype sampling is reported separately from the time for index loading and building. GraphAligner does not report the time used for indexing. Runtime includes the time for indexing and haplotype sampling.

| Reference | Mapper | True Positive | False Negative | False Positive | Recall | Precision | F1 |
|---|---|---|---|---|---|---|---|
| CHM13 | Minimap2-map-hifi | 3369762 | 17221 | 11934 | 0.9949 | 0.9965 | 0.9957 |
| CHM13 | PBMM2 | 3368694 | 18322 | 13183 | 0.9946 | 0.9961 | 0.9953 |
| HPRC-d46 | Giraffe | 3371359 | 15706 | 8763 | 0.9954 | 0.9974 | 0.9964 |
| HPRC-Sampled16 | Giraffe | 3372046 | 15016 | 8874 | 0.9956 | 0.9974 | 0.9965 |

Table S9: **DeepVariant SNP calling results with HiFi reads**.

| Reference | Mapper | True Positive | False Negative | False Positive | Recall | Precision | F1 |
|---|---|---|---|---|---|---|---|
| CHM13 | Minimap2-lr:hqae | 3343859 | 43304 | 14621 | 0.9872 | 0.9956 | 0.9914 |
| HPRC-d46 | Giraffe | 3344222 | 42955 | 12842 | 0.9873 | 0.9962 | 0.9917 |
| HPRC-Sampled16 | Giraffe | 3344942 | 42230 | 12545 | 0.9875 | 0.9963 | 0.9919 |

Table S10: **DeepVariant SNP calling results with R10 reads**.

| Reference | Mapper | True Positive | False Negative | False Positive | Recall | Precision | F1 |
|---|---|---|---|---|---|---|---|
| CHM13 | Minimap2-map-hifi | 846279 | 36734 | 25958 | 0.9584 | 0.9695 | 0.9639 |
| CHM13 | PBMM2 | 845528 | 37497 | 28492 | 0.9575 | 0.9668 | 0.9621 |
| HPRC-d46 | Giraffe | 846568 | 36466 | 26010 | 0.9587 | 0.9696 | 0.9641 |
| HPRC-Sampled16 | Giraffe | 847219 | 35811 | 25597 | 0.9594 | 0.9701 | 0.9647 |

Table S11: **DeepVariant indel calling results with HiFi reads**.

| Reference | Mapper | True Positive | False Negative | False Positive | Recall | Precision | F1 |
|---|---|---|---|---|---|---|---|
| CHM13 | Minimap2-lr:hqae | 637743 | 245324 | 74895 | 0.7222 | 0.8931 | 0.7986 |
| HPRC-d46 | Giraffe | 645419 | 237650 | 72425 | 0.7309 | 0.8973 | 0.8056 |
| HPRC-Sampled16 | Giraffe | 646287 | 236780 | 72288 | 0.7319 | 0.8976 | 0.8063 |

Table S12: **DeepVariant indel calling results with R10 reads**.

| Reference | Mapper | Genotyper | True Positive | False Negative | False Positive | Recall | Precision | F1 |
|---|---|---|---|---|---|---|---|---|
| CHM13 | Minimap2-map-hifi | Sniffles2 | 17645 | 1159 | 133 | 0.9384 | 0.9920 | 0.9645 |
| CHM13 | PBMM2 | Sniffles2 | 17483 | 1338 | 122 | 0.9289 | 0.9926 | 0.9597 |
| HPRC-d46 | Giraffe | vg call | 16153 | 2106 | 612 | 0.8847 | 0.9627 | 0.9220 |
| HPRC-d46 | GraphAligner-fast | vg call | 16218 | 2080 | 673 | 0.8863 | 0.9594 | 0.9214 |
| HPRC-Sampled16 | Giraffe | vg call | 18376 | 790 | 704 | 0.9588 | 0.9627 | 0.9607 |

Table S13: **Structural variant calling/genotpying results with HiFi reads**. Structural variants were called using Sniffles2 with Minimap2 using various presets. Structural variants were genotyped on the HPRC-d46 graph using `vg call` using Giraffe and GraphAligner, and on the HPRC-Sampled16 graph with Giraffe.



| Reference | Mapper | Genotyper | True Positive | False Negative | False Positive | Recall | Precision | F1 |
|---|---|---|---|---|---|---|---|---|
| CHM13 | Minimap2-lr:hqae | Sniffles2 | 17068 | 1718 | 133 | 0.9085 | 0.9918 | 0.9484 |
| HPRC-d46 | Giraffe | vg call | 16202 | 2086 | 615 | 0.8859 | 0.9627 | 0.9227 |
| HPRC-d46 | GraphAligner-fast | vg call | 16255 | 2056 | 728 | 0.8877 | 0.9563 | 0.9207 |
| HPRC-Sampled16 | Giraffe | vg call | 18569 | 779 | 859 | 0.9597 | 0.9553 | 0.9575 |

Table S14: **Structural variant calling/genotyping results with R10 reads**. Structural variants were called using Sniffles2 with Minimap2 using various presets. Structural variants were genotyped on the HPRC-d46 graph using `vg call` using Giraffe and GraphAligner, and on the HPRC-Sampled16 graph with Giraffe.

#### S8 Supplemental Figures

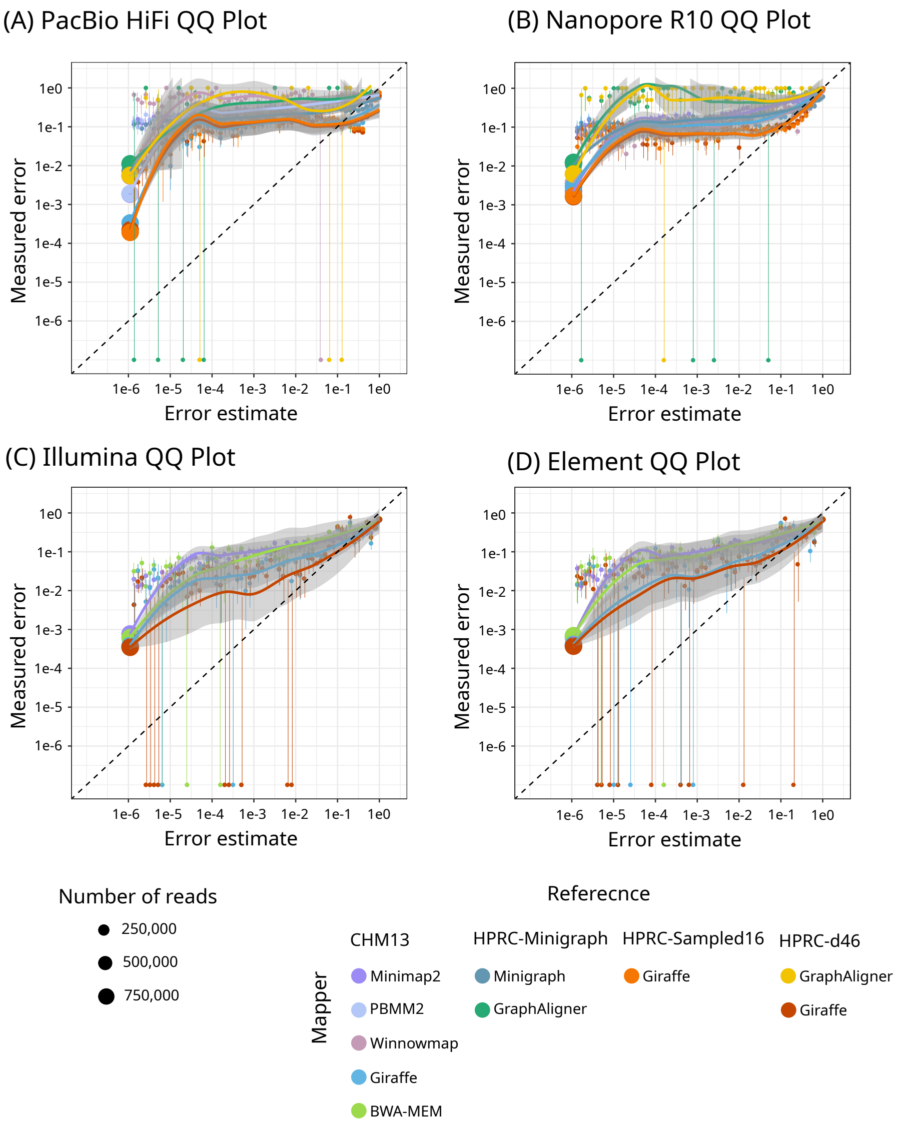


Figure S4: **Long and short read QQ plots**. QQ plots for 1 million simulated HiFi (A), R10 (B), Illumina (C), and Element (D) reads.

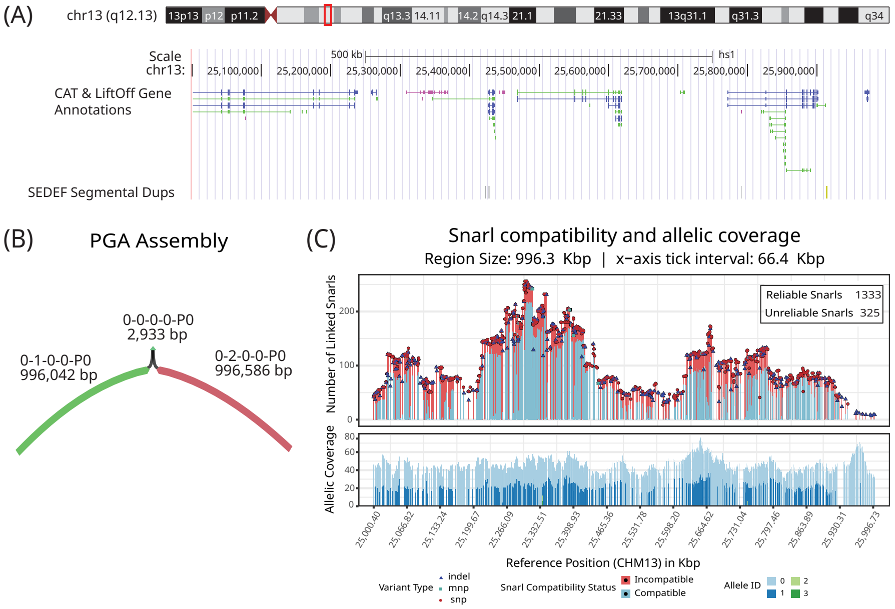


(D) PGA vs. Hifiasm : Assembly Alignment to HG002 Reference Statistics

| Assembler | Segment Name | Length (bps) | Unphased/ Phased | Parental Haplotype | Mismatches | Insertions (bps) total | Insertions short indels | Insertions long indels (>10 bp) [count * indel_length] | Deletions (bps) total | Deletions short indels | Deletions long indels (>10 bp) [count * indel_length] |
|---|---|---|---|---|---|---|---|---|---|---|---|
| PGA | 0-2-0-0-P0 | 996,805 | P | PAT | 0 | 33 | 33 | 0 | 219 | 119 | 1*20 1*80 |
| | 0-0-0-0-P0 | 2,933 | U | PAT | 0 | 0 | 0 | 0 | 0 | 0 | 0 |
| | 0-1-0-0-P0 | 996,448 | P | MAT | 0 | 36 | 36 | 0 | 406 | 100 | 1*10 1*296 |
| | 0-0-0-0-P0 | 2,933 | U | MAT | 0 | 0 | 0 | 0 | 0 | 0 | 0 |
| Hifiasm | utg028023l | 999,759 | P | PAT | 0 | 53 | 53 | 0 | 95 | 95 | 0 |
| | utg028024l | 999,394 | P | MAT | 0 | 48 | 48 | 0 | 89 | 89 | 0 |

Figure S5: **PGA assembly of a 1 Mbp region on chromosome 13**. (A) UCSC Genome Browser view of the ROI (chr13:25,000,000–26,000,000) on T2T-CHM13v2.0, showing CAT gene annotations and the SEDEF SD track. (B) Bandage view of the PGA assembly of the ROI, with segments showing Shasta contigs labeled by ID and length. (C) Snarl-compatibility and allelic-coverage plot summarizing phasing consistency across snarls. The x-axis is the position of each snarl along the CHM13 path in the graph. On the top panel, the bars represent linked snarls colored by compatibility status. Overlaid symbols indicate variant type (SNP, MNP, or indel). The lower panel displays per-allele coverage for reliable snarls. 1333 of 1658 snarls were found to be reliable and used for generating anchors. (D) Comparison of PGA and Hifiasm assemblies aligned to HG002 haplotypes: The table compares both assemblies in terms of contig lengths, counts of mismatches (SNPs) and indels (by size), and assembly contiguity.

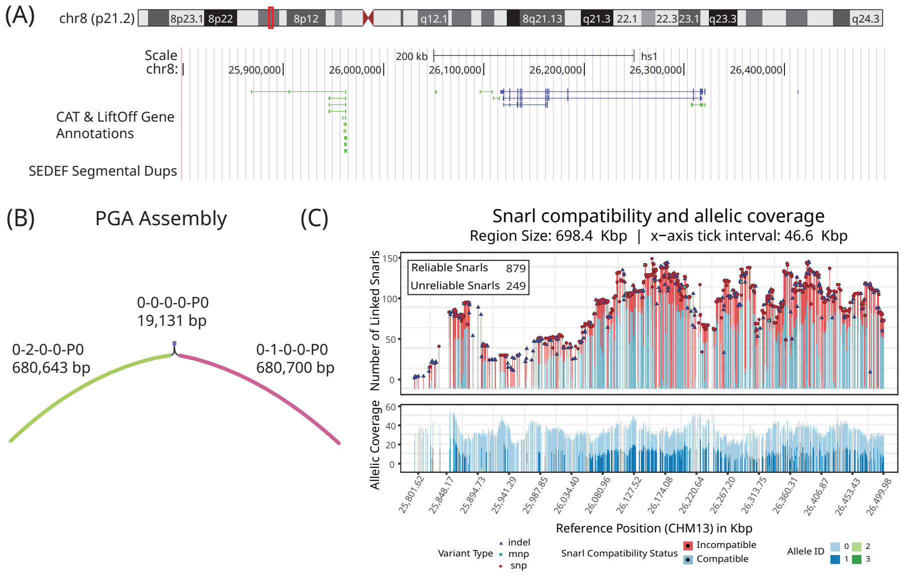


(D) PGA vs. Hifiasm : Assembly Alignment to HG002 Reference Statistics

| Assembler | Segment Name | Length (bps) | Unphased/ Phased | Parental Haplotype | Mismatches | Insertions (bps) | | | Deletions (bps) | | |
|---|---|---|---|---|---|---|---|---|---|---|---|
| | | | | | | total | short indels | long indels (>10 bp) [count * indel_length] | total | short indels | long indels (>10 bp) [count * indel_length] |
| PGA | 0-0-0-0-P0 | 19,134 | U | PAT | 1 | 1 | 1 | 0 | 3 | 3 | 0 |
| | 0-1-0-0-P0 | 680,824 | P | PAT | 0 | 22 | 22 | 0 | 123 | 107 | 1*16 |
| | 0-0-0-0-P0 | 19,132 | U | MAT | 0 | 0 | 0 | 0 | 1 | 1 | 0 |
| | 0-2-0-0-P0 | 680,907 | P | MAT | 0 | 22 | 22 | 0 | 264 | 104 | 1*16<br>1*18<br>1*26<br>1*37<br>1*63 |
| Hifiasm | utg019344l | 62,947 | P | PAT | 0 | 1 | 1 | 0 | 5 | 5 | 0 |
| | utg019347l | 37,877 | P | PAT | 1 | 4 | 4 | 0 | 4 | 4 | 0 |
| | utg019348l | 145,997 | P | PAT | 0 | 12 | 12 | 0 | 12 | 12 | 0 |
| | utg019349l | 560,563 | P | PAT | 2 | 30 | 30 | 0 | 77 | 77 | 0 |
| | utg019345l | 33,251 | P | MAT | 0 | 1 | 1 | 0 | 4 | 4 | 0 |
| | utg019346l | 179,098 | P | MAT | 0 | 8 | 8 | 0 | 25 | 25 | 0 |
| | utg019350l | 49,448 | P | MAT | 0 | 1 | 1 | 0 | 8 | 8 | 0 |
| | utg019351l | 532,620 | P | MAT | 0 | 26 | 26 | 0 | 68 | 68 | 0 |

Figure S6: **PGA assembly of a 700Kb region on chromosome 8**. (A) UCSC Genome Browser view of the ROI (chr8:25,800,000-26,500,000) on T2T-CHM13v2.0, showing CAT gene annotations and the SEDEF SD track. (B)Bandage view of the PGA assembly of the ROI, with segments showing Shasta contigs labeled by ID and length. (C) Snarl-compatibility and allelic-coverage plot summarizing phasing consistency across snarls. The x-axis is the position of each snarl along the CHM13 path in the graph. On the top panel, the bars represent linked snarls colored by compatibility status. Overlaid symbols indicate variant type (SNP, MNP, or indel). The lower panel displays per-allele coverage for reliable snarls. 879 of 1128 snarls were found to be reliable and used for generating anchors. (D) Comparison of PGA and Hifiasm assemblies aligned to HG002 haplotypes: The table compares both assemblies in terms of contig lengths, counts of mismatches (SNPs) and indels (by size), and assembly contiguity.

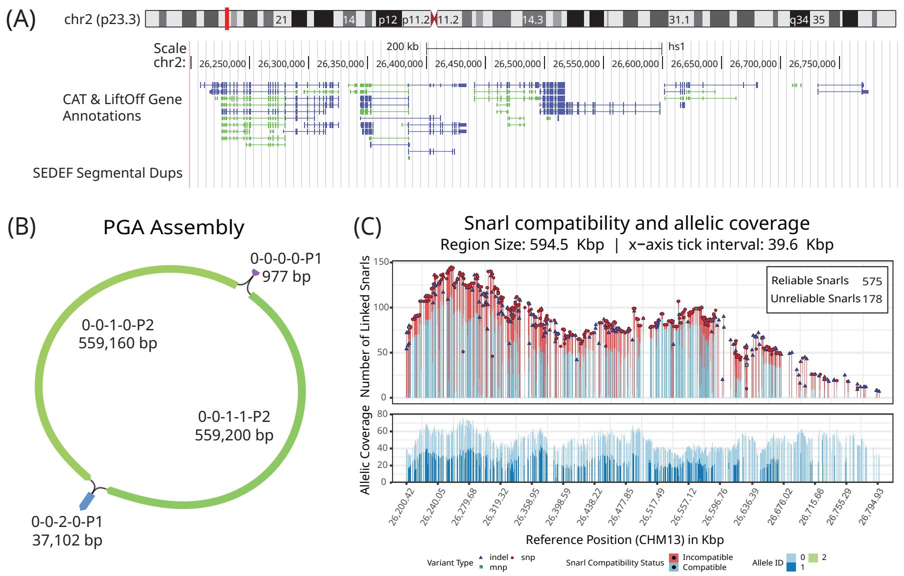


(D) PGA vs. Hifiasm : Assembly Alignment to HG002 Reference Statistics

| Assembler | Segment Name | Length (bps) | Unphased/ Phased | Parental Haplotype | Mismatches | Insertions (bps) total | Insertions short indels | Insertions long indels (>10 bp) [count * indel_length] | Deletions (bps) total | Deletions short indels | Deletions long indels (>10 bp) [count * indel_length] |
|---|---|---|---|---|---|---|---|---|---|---|---|
| PGA | 0-0-0-0-P1 | 977 | U | PAT | 0 | 2 | 2 | 0 | 0 | 0 | 0 |
| | 0-0-1-0-P2 | 559,237 | P | PAT | 0 | 18 | 18 | 0 | 77 | 59 | 1*18 |
| | 0-0-2-0-P1 | 37,109 | U | PAT | 0 | 4 | 4 | 0 | 7 | 7 | 0 |
| | 0-0-0-0-P1 | 979 | U | MAT | 0 | 2 | 2 | 0 | 2 | 2 | 0 |
| | 0-0-1-1-P2 | 559,264 | P | MAT | 0 | 27 | 27 | 0 | 64 | 64 | 0 |
| | 0-0-2-0-P1 | 37,109 | U | MAT | 0 | 4 | 4 | 0 | 7 | 7 | 0 |
| Hifiasm | utg003807l | 560,143 | P | PAT | 0 | 24 | 24 | 0 | 63 | 63 | 0 |
| | utg003809l | 19,546 | P | PAT | 1 | 8 | 8 | 0 | 1 | 1 | 0 |
| | utg003810l | 61,728 | P | PAT | 0 | 5 | 5 | 0 | 1 | 1 | 0 |
| | utg003806l | 523,873 | P | MAT | 0 | 33 | 33 | 0 | 56 | 56 | 0 |
| | utg003808l | 68,198 | P | MAT | 0 | 2 | 2 | 0 | 2 | 2 | 0 |
| | utg003811l | 48,454 | P | MAT | 0 | 6 | 6 | 0 | 2 | 2 | 0 |
| | utg003812l | 42,795 | P | MAT | 0 | 5 | 5 | 0 | 4 | 4 | 0 |

Figure S7: **PGA assembly of a 600Kb region on chromosome 2**. (A) UCSC Genome Browser view of the ROI (chr2:26,200,000-26,800,000) on T2T-CHM13v2.0, showing CAT gene annotations and the SEDEF SD track. (B) Bandage view of the PGA assembly of the ROI, with segments showing Shasta contigs labeled by ID and length. (C) Snarl-compatibility and allelic-coverage plot summarizing phasing consistency across snarls. The x-axis is the position of each snarl along the CHM13 path in the graph. On the top panel, the bars represent linked snarls colored by compatibility status. Overlaid symbols indicate variant type (SNP, MNP, or indel). The lower panel displays per-allele coverage for reliable snarls. 575 of 753 snarls were found to be reliable and used for generating anchors. (D) Comparison of PGA and Hifiasm assemblies aligned to HG002 haplotypes: The table compares both assemblies in terms of contig lengths, counts of mismatches (SNPs) and indels (by size), and assembly contiguity.

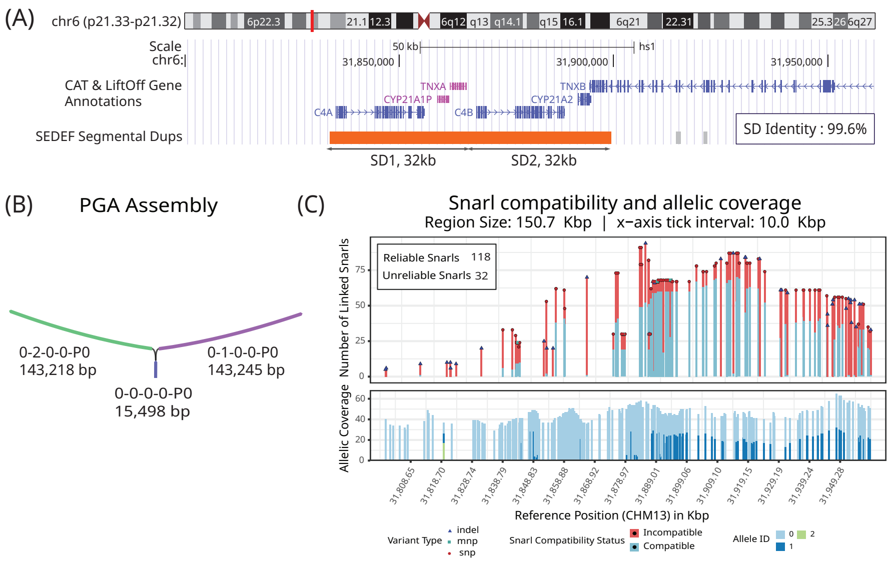


(D) PGA vs. Hifiasm : Assembly Alignment to HG002 Reference Statistics

| Assembler | Segment Name | Length (bps) | Unphased/ Phased | Parental Haplotype | Mismatches | Insertions (bps) | | | Deletions (bps) | | |
|---|---|---|---|---|---|---|---|---|---|---|---|
| | | | | | | total | short indels | long indels (>10 bp) [count * indel_length] | total | short indels | long indels (>10 bp) [count * indel_length] |
| PGA | 0-2-0-0-P0 | 143,272 | P | PAT | 0 | 4 | 4 | 0 | 58 | 14 | 1*44 |
| | 0-0-0-0-P0 | 15,499 | U | PAT | 0 | 1 | 1 | 0 | 2 | 2 | 0 |
| | 0-1-0-0-P0 | 143,272 | P | MAT | 0 | 2 | 2 | 0 | 11 | 11 | 0 |
| | 0-0-0-0-P0 | 15,499 | U | MAT | 0 | 1 | 1 | 0 | 3 | 3 | 0 |
| Hifiasm | utg014993l | 43,605 | P | PAT | 0 | 0 | 0 | 0 | 3 | 3 | 0 |
| | utg014994l | 48,070 | P | PAT | 0 | 0 | 0 | 0 | 3 | 3 | 0 |
| | utg014996l | 145,210 | P | PAT | 0 | 0 | 0 | 0 | 10 | 10 | 0 |
| | utg014995l | 99,506 | P | MAT | 0 | 0 | 0 | 0 | 4 | 4 | 0 |
| | utg014997l | 45,310 | P | MAT | 0 | 0 | 0 | 0 | 0 | 0 | 0 |
| | utg014998l | 78,315 | P | MAT | 0 | 0 | 0 | 0 | 8 | 8 | 0 |

Figure S8: **PGA assembly of a 162Kb region comprising the C4, CYP21 and TNX genes and pseudogenes**. (A) UCSC Genome Browser view of the ROI (chr6:31,798,842-31,960,826) on T2T-CHM13v2.0, showing CAT gene annotations and the SEDEF SD track. The region contains $\tilde{3}$0kb tandemly duplicated SD modules, (99.6% identity) known as the **RCCX locus**, harboring genes where copy-number variations are linked to Congenital Adrenal Hyperplasia (*CYP21A2*), Ehlers-Danlos syndrome (*TNXB*), and autoimmune-related disorders (*C4A*). (B) Bandage view of the PGA assembly of the ROI, with segments showing Shasta contigs labeled by ID and length. (C) Snarl-compatibility and allelic-coverage plot summarizing phasing consistency across snarls. The x-axis is the position of each snarl along the CHM13 path in the graph. On the top panel, the bars represent linked snarls colored by compatibility status. Overlaid symbols indicate variant type (SNP, MNP, or indel). The lower panel displays per-allele coverage for reliable snarls. 118 of 150 snarls were found to be reliable and used for generating anchors. (D) Comparison of PGA and Hifiasm assemblies aligned to HG002 haplotypes: The table compares both assemblies in terms of contig lengths, counts of mismatches (SNPs) and indels (by size), and assembly contiguity.

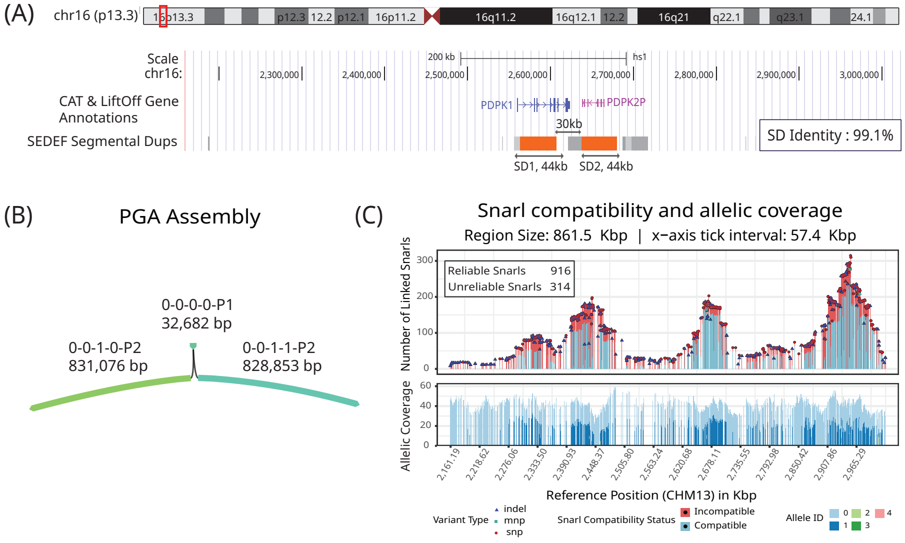


(D) PGA vs. Hifiasm : Assembly Alignment to HG002 Reference Statistics

| Assembler | Segment Name | Length (bps) | Unphased/ Phased | Parental Haplotype | Mismatches | Insertions (bps) | | | Deletions (bps) | | |
|---|---|---|---|---|---|---|---|---|---|---|---|
| | | | | | | total | short indels | long indels (>10 bp) [count * indel_length] | total | short indels | long indels (>10 bp) [count * indel_length] |
| PGA | 0-0-0-0-P1 | 25,580 | U | PAT | 0 | 0 | 0 | 0 | 9 | 9 | 0 |
| | 0-0-1-0-P2 | 837,304 | P | PAT | 0 | 69 | 69 | 0 | 157 | 115 | 1*14 1*28 |
| | 0-0-0-0-P1 | 25,580 | U | MAT | 0 | 0 | 0 | 0 | 13 | 13 | 0 |
| | 0-0-1-1-P2 | 834,967 | P | MAT | 0 | 62 | 62 | 0 | 132 | 132 | 0 |
| Hifiasm | utg031383l | 59,686 | P | PAT | 0 | 1 | 1 | 0 | 6 | 6 | 0 |
| | utg031385l | 997,569 | P | PAT | 0 | 82 | 82 | 0 | 107 | 107 | 0 |
| | utg031384l | 1,025,419 | P | MAT | 0 | 74 | 74 | 0 | 122 | 119 | 1*12 |

Figure S9: **PGA assembly of a 862Kb region comprising the *PDPK1* gene and *PDPK2P* pseudogene**. (A) UCSC Genome Browser view of the ROI (chr16:2,160,175-3,022,916) on T2T-CHM13v2.0, showing CAT gene annotations (one representative transcript per gene shown) and the SEDEF SD track. *PDPK1* and *PDPK2P* (a lncRNA) promote tumor progression by activating the PI3K/AKT pathway, making them promising therapeutic targets. They reside within SDs, sharing 99.1% sequence identity. (B) Bandage view of the PGA assembly of the ROI, with segments showing Shasta contigs labeled by ID and length. (C) Snarl-compatibility and allelic-coverage plot summarizing phasing consistency across snarls. The x-axis is the position of each snarl along the CHM13 path in the graph. On the top panel, the bars represent linked snarls colored by compatibility status. Overlaid symbols indicate variant type (SNP, MNP, or indel). The lower panel displays per-allele coverage for reliable snarls. 916 of 1230 snarls were found to be reliable and used for generating anchors. (D) Comparison of PGA and Hifiasm assemblies aligned to HG002 haplotypes: The table compares both assemblies in terms of contig lengths, counts of mismatches (SNPs) and indels (by size), and assembly contiguity.

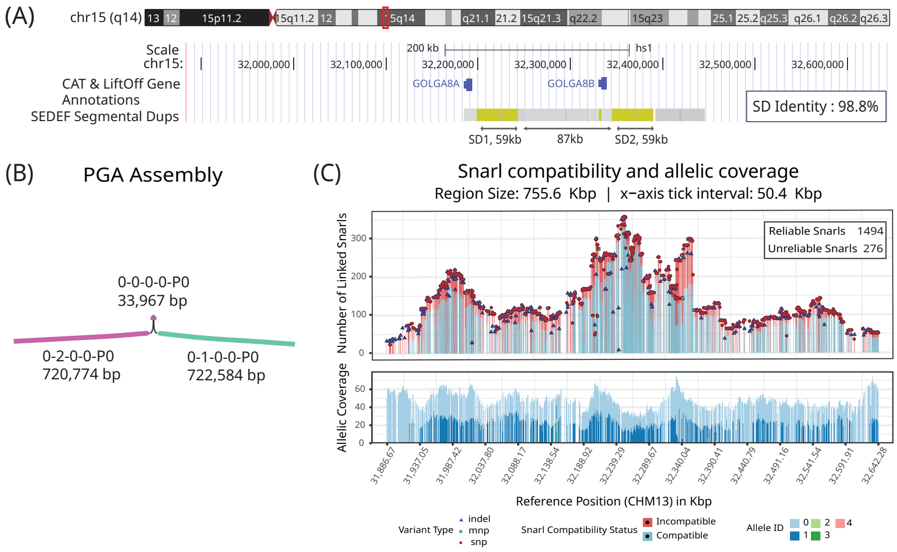


(D) PGA vs. Hifiasm : Assembly Alignment to HG002 Reference Statistics

| Assembler | Segment Name | Length (bps) | Unphased/ Phased | Parental Haplotype | Mismatches | Insertions (bps) | | | Deletions (bps) | | |
|---|---|---|---|---|---|---|---|---|---|---|---|
| | | | | | | total | short indels | long indels (>10 bp) [count * indel_length] | total | short indels | long indels (>10 bp) [count * indel_length] |
| PGA | 0-0-0-0-P0 | 33,974 | U | PAT | 0 | 1 | 1 | 0 | 7 | 7 | 0 |
| | 0-1-0-0-P0 | 720,898 | P | PAT | 0 | 32 | 32 | 0 | 124 | 124 | 0 |
| | 0-0-0-0-P0 | 33,977 | U | MAT | 1 | 1 | 1 | 0 | 10 | 10 | 0 |
| | 0-2-0-0-P0 | 722,929 | P | MAT | 0 | 19 | 19 | 0 | 345 | 345 | 0 |
| Hifiasm | utg030399l | 754,782 | P | PAT | 0 | 38 | 38 | 0 | 96 | 96 | 0 |
| | utg030398l | 756,834 | P | MAT | 0 | 35 | 35 | 0 | 88 | 88 | 0 |

Figure S10: **PGA assembly of a 758Kb region comprising the *GOLGA8A* and *GOLGA8B* genes**. (A) UCSC Genome Browser view of the ROI (chr15:31,884,805-32,643,157) on T2T-CHM13v2.0, showing CAT gene annotations (one representative transcript per gene shown) and the SEDEF SD track. *GOLGA8A* and *GOLGA8B* are paralogous golgin proteins embedded within highly similar SDs (98.8% identity). Their repetitive nature creates instability hotspots linked to chromosomal deletions underlying neurodevelopmental disorders (intellectual disability, autism, epilepsy, schizophrenia), while *GOLGA8B* has also been associated with cancer invasion and metastasis. (B) Bandage view of the PGA assembly of the ROI, with segments showing Shasta contigs labeled by ID and length. (C) Snarl-compatibility and allelic-coverage plot summarizing phasing consistency across snarls. The x-axis is the position of each snarl along the CHM13 path in the graph. On the top panel, the bars represent linked snarls colored by compatibility status. Overlaid symbols indicate variant type (SNP, MNP, or indel). The lower panel displays per-allele coverage for reliable snarls. 1494 of 1770 snarls were found to be reliable and used for generating anchors. (D) Comparison of PGA and Hifiasm assemblies aligned to HG002 haplotypes: The table compares both assemblies in terms of contig lengths, counts of mismatches (SNPs) and indels (by size), and assembly contiguity.
